# Wnt-regulated lncRNA discovery enhanced by *in vivo* identification and CRISPRi functional validation

**DOI:** 10.1101/2020.02.09.941005

**Authors:** Shiyang Liu, Nathan Harmston, Trudy Lee Glaser, Yunka Wong, Zheng Zhong, Babita Madan, David M. Virshup, Enrico Petretto

**Affiliations:** Program in Cancer and Stem Cell Biology, Duke-NUS Medical School, 8 College Rd, Singapore 169857, Singapore; Program in Cardiovascular and Metabolic Disorders, Duke-NUS Medical School, 8 College Rd, Singapore 169857, Singapore; Science Division, Yale-NUS College, 16 College Avenue West #01-220, 138527, Singapore; Department of Pediatrics, Duke University School of Medicine, 128 Davison Bldg, Durham, NC 27710, USA; MRC London Institute of Medical Sciences, Imperial College London, Du Cane Rd, London W12 0NN, United Kingdom.

**Keywords:** functional lncRNAs, Wnt signaling, cancer, CRISPRi screen

## Abstract

**Background:** Wnt signaling is an evolutionarily conserved developmental pathway that is frequently hyperactivated in cancer. While multiple protein-coding genes regulated by Wnt signaling are known, the functional lncRNAs regulated by Wnt signaling have not been systematically characterized.

**Results:** We comprehensively mapped lncRNAs from an orthotopic Wnt-addicted pancreatic cancer model, identifying 3,633 lncRNAs, of which 1,503 were regulated by Wnt signaling. We found lncRNAs were much more sensitive to changes in Wnt signaling in xenografts than in cultured cells. To functionally validate Wnt-regulated lncRNAs, we performed CRISPRi screens to assess their role in cancer cell proliferation. Consistent with previous genome-wide lncRNA CRISPRi screens, around 1% (13/1,503) of the Wnt-regulated lncRNAs could modify cancer cell growth *in vitro*. This included *CCAT1* and *LINC00263*, previously reported to regulate cancer growth. Using an *in vivo* CRISPRi screen, we doubled the discovery rate, identifying twice as many Wnt-regulated lncRNAs (25/1,503) that had a functional effect on cancer cell growth.

**Conclusions:** Our study demonstrates the value of studying lncRNA functions *in vivo*, provides a valuable resource of lncRNAs regulated by Wnt signaling and establishes a framework for systematic discovery of functional lncRNAs.

## Background

lncRNAs play key roles in diverse biological processes, ranging from development, such as *XIST* for dosage compensation (Brown et al., 1991) and *H19* for imprinting (Brannan et al., 1990), to different diseases including cancer (Huarte, 2015). lncRNAs have been shown to play important roles in fundamental biological signaling pathways regulated by P53, Notch and TGF-β (Huarte et al., 2010; Trimarchi et al., 2014; Yuan et al., 2014). lncRNAs can contribute to the development of cancer through aberrant expression or mutation, altering their normal physiological functions in signaling pathways (Schmitt & Chang, 2016). Advancements in transcriptomics have greatly expanded the number of long noncoding RNAs (lncRNAs) annotated in the human genome (Hon et al., 2017; Iyer et al., 2015), but only a small fraction have been characterized at a functional level.

Wnt/β-catenin signaling is an important evolutionarily conserved signaling pathway β that is crucial for embryonic development and tissue regeneration (Nusse & Clevers, 2017). After Wnt ligands binding to Frizzled and other co-receptors on the cell surface, β-catenin is stabilized and translocates into the nucleus, where it interacts with TCF/LEF transcription factors in a context-dependent manner to regulate the expression of multiple protein-coding genes such as *MYC* and *AXIN2*. Dysregulation of Wnt signalling is found in multiple cancers. The most common mutations activating Wnt/β-catenin signaling occur in colorectal cancer, where truncations of APC cause abnormal stabilization of β-catenin and constitutive transcriptional activation (Polakis, 2012; Zhan et al., 2017; Zhong & Virshup, 2019). A different class of mutations confer cancer dependency on Wnt ligands. For example, *RNF43 and RPSO3* mutations cause increased abundance of Wnt receptors on the cell surface, making the cancer cells addicted to Wnt signaling (X. Jiang et al., 2013; Koo et al., 2012; Seshagiri et al., 2012). RNF43 mutations are found in 5 – 10% of pancreatic cancers, while RPSO3 translocations are found in 10% of colorectal cancers (Bailey et al., 2016; Cancer Genome Atlas Research Network, 2017; Giannakis et al., 2014; Seshagiri et al., 2012; Waddell et al., 2015).

Wnt addiction in cancer presents a therapeutic opportunity (Madan & Virshup, 2015). All Wnts require palmitoleation in the endoplasmic reticulum by the enzyme PORCN for their secretion and function (Willert et al., 2003). Small molecule PORCN inhibitors block this modification and hence the activity of all Wnts. We and others have demonstrated that PORCN inhibitors such as ETC-159 suppress the growth of Wnt-addicted cancers in multiple preclinical models (B. Chen et al., 2009; X. Jiang et al., 2013; Madan et al., 2016). Due to its efficacy, the PORCN inhibitor ETC-159 has advanced to clinical trials (Ng et al., 2017). ETC-159 is also a useful research tool to study Wnt dependent genes. We found that more than 75% of the transcriptome responded to PORCN inhibition by ETC-159 in Wnt-addicted cancers, with significantly more genes changing *in vivo* than *in vitro* (Madan et al., 2018, 2016). Thus, PORCN inhibition is a powerful tool to study Wnt-regulated genes, and these Wnt-regulated genes are best studied *in vivo* in the presence of the appropriate microenvironment.

To date, only a few individual lncRNAs have been linked to Wnt signaling. For example, *MYU* (*VPS9D1-AS1*) is a target of Wnt/c-Myc signaling involved in colon cancer (Kawasaki et al., 2016). However, currently there are no systematic studies on functional lncRNAs regulated by Wnt signaling *in vivo*. Here, we comprehensively mapped Wnt-regulated lncRNAs from an orthotopic Wnt-addicted pancreatic cancer model and determined their wider roles in other cancers. To functionally validate the Wnt-regulated lncRNAs, we performed CRISPRi screens both *in vitro* and *in vivo*. Notably, we found multiple Wnt-regulated lncRNAs that had functional effects on cancer cell growth only in a xenograft model, demonstrating the value of studying lncRNA functions *in vivo*. This study provides a valuable resource of functional lncRNAs regulated by Wnt signaling. It also establishes a framework that can be broadly adapted for systematic discovery and functional annotation and validation of lncRNAs *in vivo*.

## Results

### Discovery of Wnt-regulated lncRNAs

The HPAF-II pancreatic cancer cells contain a *RNF43* missense mutation that makes them addicted to Wnt signaling. As previously reported, mice with established orthotopic HPAF-II xenografts were treated with the PORCN inhibitor ETC-159 for 7 days. Tumors were harvested for transcriptomic analysis at indicated time points (0, 3, 8, 16, 32, 56 and 168 hours) after starting ETC-159 treatment. The data were previously analyzed with a focus on protein-coding genes and splice variants (Idris et al., 2019; Madan et al., 2018). To comprehensively identify Wnt-regulated lncRNAs in pancreatic cancer *in vivo*, we reanalyzed this time-course transcriptomic dataset (Figure 1A). We first used *de novo* assembly to comprehensively identify all the putative transcripts in this Wnt-addicted pancreatic xenograft model. These transcripts were then compared with the Ensembl build 79 transcriptome to identify putative novel lncRNAs. The putative novel lncRNAs were filtered based on their length (> 200 bp), and we eliminated those with coding potential called by any of three computational tools: CPAT (L. Wang et al., 2013), CPC (Kong et al., 2007) and Slncky (J. Chen et al., 2016) (see Methods for details). The novel lncRNAs were combined with previously annotated lncRNAs from Ensembl build 79 to establish a comprehensive list of lncRNAs present in our RNA-seq dataset. We next selected all the lncRNA genes with TPM > 1. Using these stringent criteria, we identified a set of 3,633 lncRNAs in an orthotopic *RNF43*-mutant pancreatic cancer model (Figure 1A). Amongst these 3,633 lncRNAs, we found that the expression of 1,503 lncRNAs changed over time upon Wnt inhibition (false discovery rate (FDR) < 5%), therefore we refer to these lncRNAs as “Wnt-regulated lncRNAs” (Table S1). Among the 1,503 Wnt-regulated lncRNAs, 325 lncRNAs were not annotated in Ensembl build 79. We further compared these novel lncRNAs with FANTOM5 lncRNA annotations (Hon et al., 2017) and found 172 lncRNAs that have not been previously annotated either in Ensembl or FANTOM5 (Figure 1B).

**Figure 1:**
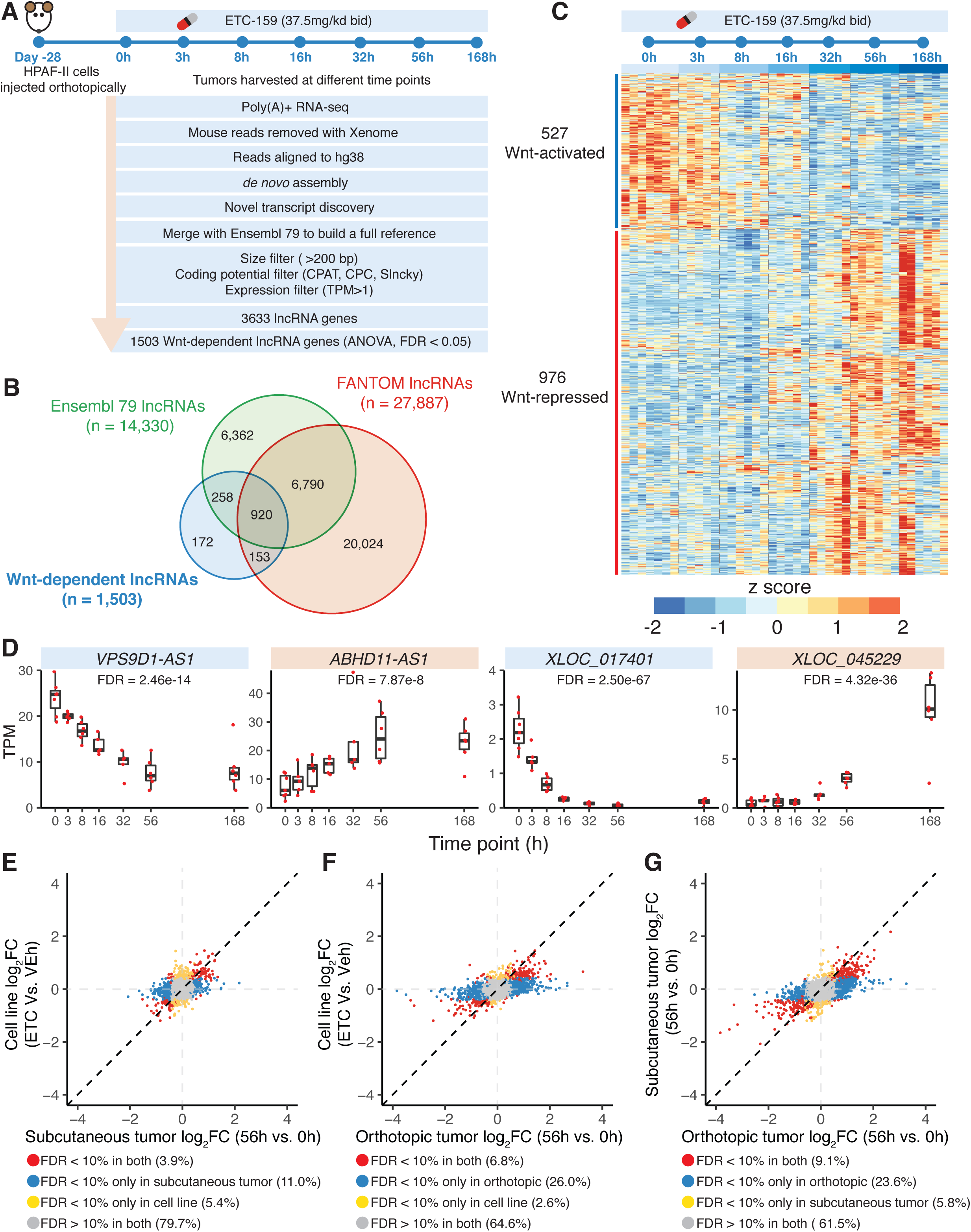
Identification of Wnt-regulated lncRNAs from orthotopic *RNF43*-mutant pancreatic cancer model. **(A)** Computational pipeline to identify 1,503 Wnt-regulated lncRNAs from orthotopic RNF43-mutant pancreatic cancer. **(B)** Comparison of Wnt-regulated lncRNAs with Ensembl build 79 and FANTOM5 lncRNA annotations. **(C)** Expression profiles of 1,503 Wnt-regulated lncRNAs across time points after Wnt inhibition. **(D)** Gene expression of selected Wnt-regulated lncRNAs, including annotated lncRNAs (*VPS9D1-AS1* and *ABHD11-AS1*) and novel lncRNAs (*XLOC_017401* and *XLOC_045229*). TPM, transcripts per million. **(E, F, G)** Fold change of lncRNAs after Wnt inhibition compared across models. More lncRNAs respond to Wnt inhibition in the HPAF-II subcutaneous **(E)** and orthotopic models **(F)** than in HPAF-II cells cultured *in vitro*. FC, fold change. **(G)** More lncRNAs respond to Wnt inhibition in HPAF-II orthotopic model than in the subcutaneous model.

We found that twice as many lncRNAs were upregulated (976 Wnt-repressed lncRNAs) than downregulated (527 Wnt-activated lncRNAs) following PORCN inhibitor treatment (Figure 1C). Among them, 240 Wnt-repressed and 85 Wnt-activated lncRNAs are not annotated in Ensembl build 79. The 527 Wnt-activated lncRNAs responded as early as 3 hours after the first dose of ETC-159, consistent with direct regulation by Wnt/β-catenin signaling. Conversely, the 976 Wnt-repressed lncRNAs responded more slowly to Wnt inhibition (Figure 1C), which could be due to indirect Wnt regulation. For example, *VPS9D1-AS1*, a previously reported target of Wnt/MYC signaling (Kawasaki et al., 2016), was down-regulated rapidly after PORCN inhibitor treatment and the inhibition was sustained for 7 days. Similarly, a previously unannotated lncRNA *XLOC_017401* was also downregulated shortly after Wnt inhibition. In contrast, *XLOC_045229*, another previously unannotated lncRNA, was upregulated after ETC-159 treatment, but the effect was only observed after 32 hours of treatment (Figure 1D). Taken together, we identified 1,503 lncRNAs whose expression is regulated either directly or indirectly by Wnt signaling *in vivo* in an *RNF43*-mutant pancreatic cancer.

Genes that are important in cancer pathogenesis can be regulated by multiple pathways. For example, the well-known proto-oncogene *MYC* can be activated by pathological Wnt signaling in Wnt-driven cancers, and also by diverse additional pathways in other cancers (Gabay et al., 2014). Similarly, we postulated that if a specific Wnt-regulated lncRNA is important in cancer, the same lncRNA might also be dysregulated by other mechanisms in other cancer types. To test this, we analyzed gene expression data from TCGA (Goldman et al., 2019), comparing tumors with their paired normal samples. We found that many Wnt-regulated lncRNAs were also dysregulated in different and Wnt-independent types of cancers (Figure S1A). For example, 1,150 of the 1,503 Wnt-regulated lncRNAs were found in lung adenocarcinoma samples (LUAD) in TCGA, and 435 of these were significantly upregulated compared to paired normal samples (Figure 1A). We also found 253 Wnt-regulated lncRNAs exclusively upregulated or downregulated across different cancer types (Table S1). For example, *VPS9D1-AS1*, a known Wnt/MYC target, was both Wnt-activated in our study and also upregulated in 11 different types of cancers (Figure S1B), consistent with its established role as a lncRNA with oncogenic function (Kawasaki et al., 2016). Together, these analyses suggest that a subset of Wnt-regulated lncRNAs can act as mediators of oncogenic processes in both Wnt-dependent and Wnt-independent cancers.

### LncRNAs respond to Wnt inhibition more robustly *in vivo*, especially in orthotopic xenograft model

Tumor microenvironment is important for tumor pathogenesis (Miller et al., 2017; Muir & Vander Heiden, 2018; Whiteside, 2008). To examine how the response of lncRNAs to Wnt inhibition is affected by the stromal microenvironment, we compared the effect of ETC-159 on lncRNAs expression in HPAF-II orthotopic or subcutaneous xenografts (*in vivo*) and in cultured cells (*in vitro*). Nearly twice as many lncRNAs responded to the PORCN inhibitor treatment in the subcutaneous xenograft (541/3,633) compared to those that responded *in vitro* (341/3,633) (Figure 1E). A further increase in the number of lncRNAs responding to Wnt inhibition was observed in the orthotopic xenografts (1,191/3,633) (Figure 1F). This is consistent with our previous observation that Wnt-regulated gene expression changes are more robust *in vivo* (Madan et al., 2018). Interestingly, between the two *in vivo* models, many more lncRNAs responded to Wnt inhibition in the orthotopic than subcutaneous xenograft (Figure 1G). This is consistent with our previous observation that the overall changes in gene expression following Wnt inhibition were most marked in the orthotopic model (Madan et al., 2018). Taken together, this indicates that *in vivo* models can substantially enhance the discovery of Wnt-regulated genes, including lncRNAs.

### A subset of Wnt-regulated lncRNAs are co-expressed with their nearest protein-coding gene in the same TAD

Most of the Wnt-regulated lncRNAs identified here have not previously been described or functionally characterized. Since lncRNAs can be important regulators of nearby genes (Engreitz et al., 2016; Gil & Ulitsky, 2019; Luo et al., 2016), we set out to explore their potential *cis* functions. If a lncRNA and its nearby protein-coding gene (PCG) are positively co-expressed after Wnt inhibition, it suggests that the lncRNA may enhance the expression of its neighbor. To test this, we analyzed the expression changes of lncRNAs and PCGs in response to PORCN inhibitor treatment. We found that on average, Wnt-regulated lncRNAs exhibited stronger co-expression with their nearest PCG after Wnt inhibition compared to their co-expression with all PCGs (Figure S2A). This stronger co-expression can be partially explained by the fact that some of the Wnt-regulated lncRNA–nearest PCG pairs are within the same topological associated domain (TAD) (Figure S2B), where they may functionally interact with each other more frequently, as previously suggested (Dixon et al., 2012). Interestingly, for these Wnt-regulated lncRNA–nearest PCG pairs encoded within the same TAD, the PCGs were significantly enriched for Gene Ontology (GO) biological processes such as organ development and cell fate specification (Figure S2C). This suggests that these highly co-expressed Wnt-regulated lncRNAs that are proximal to PCGs and co-localized within the same TAD, are likely to be involved in the same cellular processes.

### Wnt signaling affects the *cis* functional interaction between lncRNAs and protein-coding genes

Expression quantitative trait loci (eQTLs) analysis that links DNA sequence variation with changes in gene expression has been a powerful approach for understanding the functional effects of common SNPs (Consortium & GTEx Consortium, 2017). The underlying regulatory mechanisms of the eQTL SNPs on gene expression depend on the genomic functional element perturbed by the genetic variant. For example, an eQTL SNP within a lncRNA might modify its interaction with transcription factors or epigenetic modifiers, thereby altering the expression of nearby PCGs (Gao et al., 2018). SNPs within lncRNA loci that are associated with the mRNA abundance of nearby genes (<1 Mbp apart), i.e., *cis*-acting regulation, have been systematically annotated by the FANTOM5 consortium to establish lncRNA-mRNA pairs linked by these eQTL SNPs (Hon et al., 2017). This lncRNA-mRNA interaction mediated by an eQTL suggests that these lncRNAs loci might potentially regulate the expression of nearby mRNAs. The FANTOM5 dataset contains genome-wide transcriptome profiles of 1,829 samples from more than 173 human primary cell types and 174 tissues across the human body, 276 cancer cell lines and 19 time courses of cellular treatment. If the eQTL linked lncRNA-mRNA are co-expressed in FANTOM5 samples, it further suggests a functional interaction between the lncRNA and its eQTL-linked mRNA. Here, to identify Wnt-regulated lncRNAs with potential regulatory effects on nearby PCG mRNAs, we overlapped 1,503 Wnt-regulated lncRNAs with all of the lncRNA-mRNA pairs annotated by the FANTOM5 consortium. We found 1,486 lncRNA PCG mRNA (lncRNA-PCG) pairs linked by eQTL SNPs involving 602 Wnt-regulated lncRNAs. (Some of the lncRNAs were linked to multiple PCGs, Figure 2A and Table S2). Among them, 587 lncRNA-PCG pairs were also significantly co-expressed (*p* < 0.05) in FANTOM5 samples. This co-expression across FANTOM5 samples suggests a functional interaction between the Wnt-regulated lncRNA and its eQTL-linked PCG broadly across cell types.

**Figure 2:**
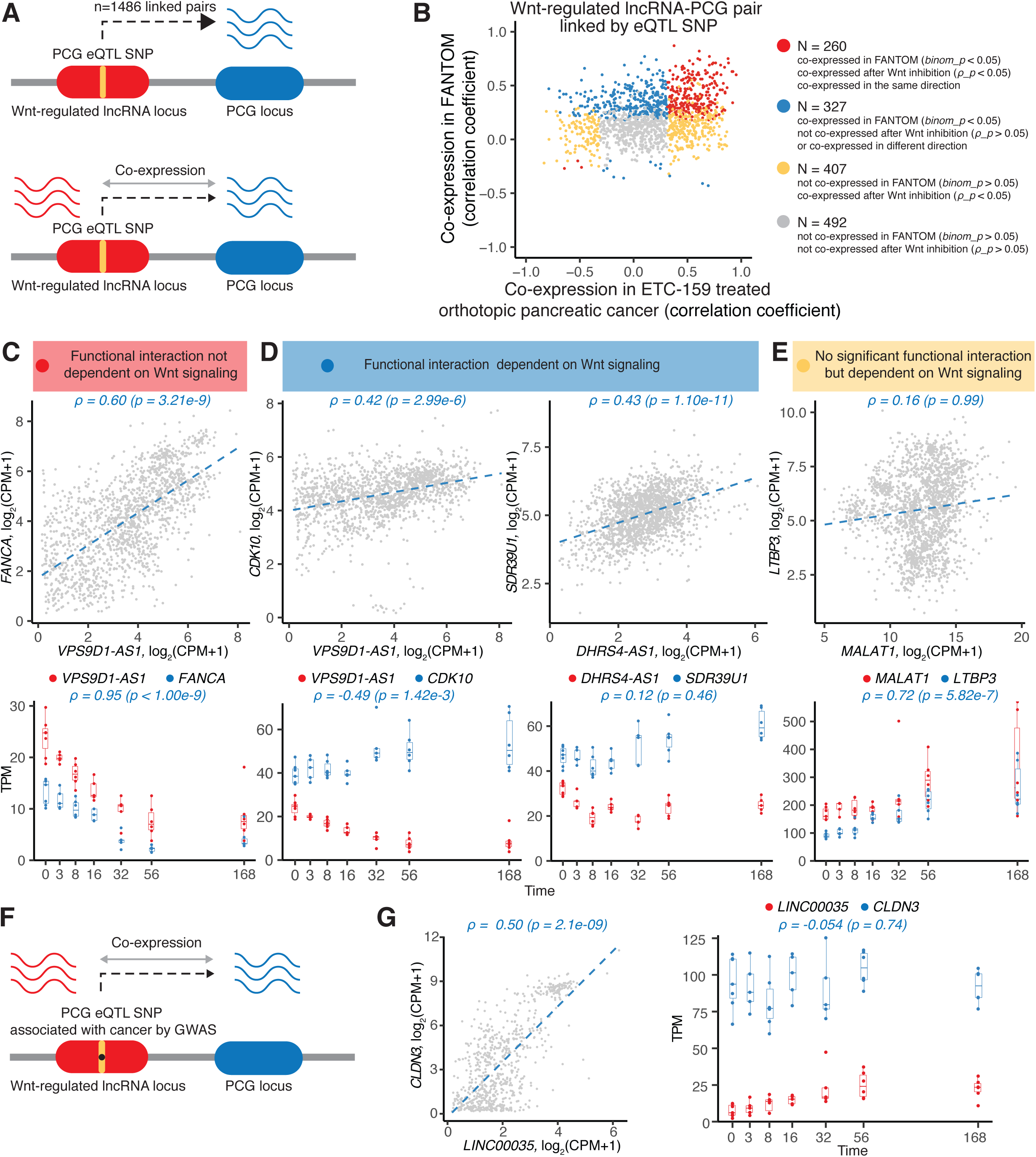
Wnt signaling affects the cis functional interaction between lncRNAs and protein-coding genes. **(A)** Wnt-regulated lncRNA is linked to its nearby protein-coding gene (PCG) if the eQTL SNP of a PCG overlaps with a lncRNA locus, as annotated by FANTOM5 consortium (Hon et al., 2017). The co-expression between Wnt-regulated lncRNA and its eQTL-linked PCG is examined both in FANTOM5 across cell types and in our dataset after Wnt inhibition. **(B)** Functional interaction between Wnt-regulated lncRNA and its eQTL-linked PCG is affected by Wnt signaling. Red, functional interaction between lncRNA-PCG pair implicated in FANTOM5 is not directly dependent on Wnt signaling; blue, functional interaction between lncRNA-PCG pair implicated in FANTOM5 is dependent on Wnt signaling; yellow, no significant functional interaction between lncRNA-PCG pair but they are co-regulated in response to Wnt-signaling; grey, lncRNA-PCG pair is neither co-regulated in response to Wnt-signaling nor functionally interacting. (C) Wnt-dependent lncRNA *VPS9D1-AS1* and its eQTL-linked PCG FANCA are functionally-interacting (*⍰* = 0.6, *p* = 3.21e-9), and the interaction is not dependent on Wnt signaling (*⍰* = 0.95, *p* < 1.00e-9). CPM, counts per million. (D) Functional interaction between Wnt-dependent lncRNA *VPS9D1-AS1* and its eQTL-linked PCG *CDK10* suggested by their co-expression (*⍰* = 0.42, *p* = 2.99e-6) across FANTOM5 samples, is dependent on Wnt signaling, as they are co-expressed in the opposite direction after Wnt inhibition (*⍰* = −0.49, *p* = 1.42e-3). Functional interaction (*⍰* = 0.43, *p* = 1.10e-11) between Wnt-dependent lncRNA *DHRS4-AS1* and its eQTL-linked PCG *SDR39U1* is also dependent on Wnt signaling, as they are not co-expressed after Wnt inhibition (*⍰* = 0.12, *p* = 0.46). (E) Wnt-dependent lncRNA *MALAT1* and its eQTL-linked PCG *LTBP3* are co-regulated in response to Wnt-signaling (*⍰* = 0.72, *p* = 5.82e-7), but not functionally-interacting (*⍰* = 0.16, *p* = 0.99). (F) 115 Wnt-regulated lncRNA-PCG pairs are linked by eQTL SNPs that are associated with cancer by GWAS. (G) Representative Wnt-regulated lncRNA *LINC0035* associated with leukemia has functional interaction with *CLDN3* (*⍰* = 0.5, *p* = 2.10e-9), and the functional interaction is dependent on Wnt signaling (*⍰* = −0.054, *p* =0.74).

We examined if Wnt signaling altered the functional interaction (co-expression) between Wnt-regulated lncRNAs and their eQTL-linked PCGs. To do this, we compared the co-expression detected in response to Wnt inhibition to the co-expression observed in the FANTOM5 dataset. First, we found 260 lncRNA-PCG pairs that were significantly co-expressed in both our dataset and FANTOM5, irrespective of Wnt signaling status (Figure 2B). One illustrative example of this consistent co-expression pattern is *VPS9D1-AS1* (lncRNA) and *FANCA* (eQTL-linked PCG) in Figure 2C. Here, the lncRNA-PCG co-expression was significant (*p* < 0.05) and had the same direction, i.e., positive in response to Wnt inhibition in our model of pancreatic cancer and positive in all FANTOM5 samples. In this set of lncRNA-PCG pairs, their functional interactions were not directly dependent on Wnt signaling. Second, there were 327 lncRNA-PCG pairs significantly co-expressed in the FANTOM5 dataset that were either not significantly co-expressed or co-expressed in the opposite direction after Wnt inhibition (Figure 2B). For example, *VPS9D1-AS1* was also linked to *CKD10* through 9 eQTL SNPs. *VPS9D1-AS1* and *CKD10* were positively co-expressed in FANTOM5 samples (⍰ = 0.42), but in response to Wnt inhibition, they were negatively co-expressed (⍰ = −0.49) (Figure 2D).

This suggests that their functional interaction is affected by Wnt signaling inhibition. Finally, a third group of 407 Wnt-regulated lncRNA-PCG pairs (Figure 2B), although linked by eQTL SNPs, were not significantly co-expressed across FANTOM5 samples. However, they were significantly co-expressed in response to Wnt inhibition in our model of pancreatic cancer. For example, *MALAT1* and *LTBP3* are not correlated in FANTOM5 but they are similarly regulated by Wnt signaling, Figure 2E. Thus, these lncRNAs and PCGs are co-regulated in a Wnt-dependent manner. Taken together, these analyses demonstrate that Wnt signaling can affect the functional interaction between Wnt-regulated lncRNAs and their eQTL-linked PCG. Therefore, Wnt signaling is important for both the regulation and the function of a subset of Wnt-regulated lncRNAs.

We then investigated the diseases associated with the Wnt-regulated lncRNA-PCG pairs linked by eQTLs, with a focus on cancer. eQTLs that co-localize with disease risk loci identified by genome-wide association studies (GWAS) are candidates for the regulation of complex traits and diseases, including SNPs associated with cancer susceptibility by GWAS (Q. Li et al., 2013). Thus we further examined the eQTL SNPs overlapping with the Wnt-regulated lncRNAs loci and matched these SNPs with those curated by FANTOM5 for 56 cancer GWAS traits. Among the 1,486 eQTL-linked Wnt-regulated lncRNA-PCG pairs, a subset of 115 pairs involving 49 Wnt-regulated lncRNAs were linked by eQTL SNPs that colocalize with cancer GWAS loci (Figure 2F, Table S3). For example, *LINC00035* was linked to *CLDN3* (Figure 2G) through 10 distinct *CLDN3* eQTL SNPs that were also associated with leukemia by GWAS (Table S3). In addition, *LINC0035* showed functional interaction with *CLDN3* in FANTOM5 (*p* = 2.10e-9), however, this functional interaction disappeared when we inhibited Wnt signaling (*p* = 0.74). This might suggest that *LINC0035* is involved in susceptibility to leukemia through its regulation of *CLDN3* in a Wnt-dependent manner. Integrating eQTL-linked Wnt-regulated lncRNA-PCG pairs with cancer GWAS data suggests that 3% (49/1,503) of the Wnt-regulated lncRNAs may confer cancer susceptibility through their *cis*-regulation of eQTL-linked PCGs (Gao et al., 2018; Tan et al., 2017).

### Wnt-regulated lncRNAs and protein-coding genes form gene networks that are dysregulated in cancers

Beside *cis* regulatory functions, lncRNAs can also participate in gene networks that regulate diverse biological processes (Guttman et al., 2011; Kopp & Mendell, 2018). To investigate which gene networks the various Wnt-regulated lncRNAs may be involved in, we performed time-series clustering on the differentially expressed Wnt-regulated lncRNAs and PCGs. This analysis closely paralleled a similar time-series clustering of PCGs that we reported previously (Madan et al., 2018). The lncRNAs and PCGs fell into 63 distinct clusters based on their pattern of expression change following Wnt inhibition (Figure S3A). The similar and coherent dynamic response of each cluster to Wnt inhibition suggests the presence of a common regulatory process within each cluster (Rotival & Petretto, 2014). As many Wnt-regulated lncRNAs and PCGs were also dysregulated in different types of cancers as determined by differential expression between tumors and their paired normal samples using the TCGA dataset (Goldman et al., 2019)(Figure S1A), we tested if the lncRNA-PCGs clusters were enriched for dysregulated genes in different cancer types. We found that 46 out of the 63 clusters were enriched (FDR < 5%) for genes dysregulated in at least one type of cancer (Figure S3B). In addition, most of these clusters (38/46) were enriched for genes consistently either up- or down-regulated in several different cancer types (Figure S3B and Figure 3A and 3B). For example, cluster 9 contained 67 Wnt-activated lncRNAs and 357 PCGs, including well established Wnt target genes (e.g., *NKD1*, *AXIN2*, *LGR5*, *MYC*, *BMP4*, *FGF9*) (Figure 3E). This cluster was enriched for genes upregulated in 6 cancer types and was significantly enriched for ncRNA metabolic process, Wnt signaling and cell differentiation. Many of the genes associated with ncRNA metabolic process (*NOP56*, *METTL1*, *RRP1*, *AIMP2*, *EXOSC5*) were also overexpressed in multiple cancers. With a few notable exceptions such as lncRNA *LINC00511* (J. Zhang et al., 2019), most of the lncRNAs in this cluster do not have established biological functions. One the other hand, cluster 2 contained mainly Wnt-repressed genes, the majority of which were downregulated in eight cancer types. The PCGs from this cluster were enriched for processes related to vesicle organization, vesicle transport and immune response. This last finding is consistent with recent studies demonstrating that Wnt signaling prevents anti-tumor immunity and suppresses immune surveillance (Holtzhausen et al., 2015; Spranger et al., 2015). Although most of the lncRNAs from cluster 2 have not been characterized before, *LINC00910* was previously identified as a lncRNA highly connected to other gene promoter regions and was proposed to be involved in lymphocyte activation (Cai et al., 2016). Taken together, this lncRNA-PCG network analysis suggests specific Wnt-regulated lncRNAs in gene networks are involved in distinct biological processes that contribute to the pathogenesis of cancers.

**Figure 3:**
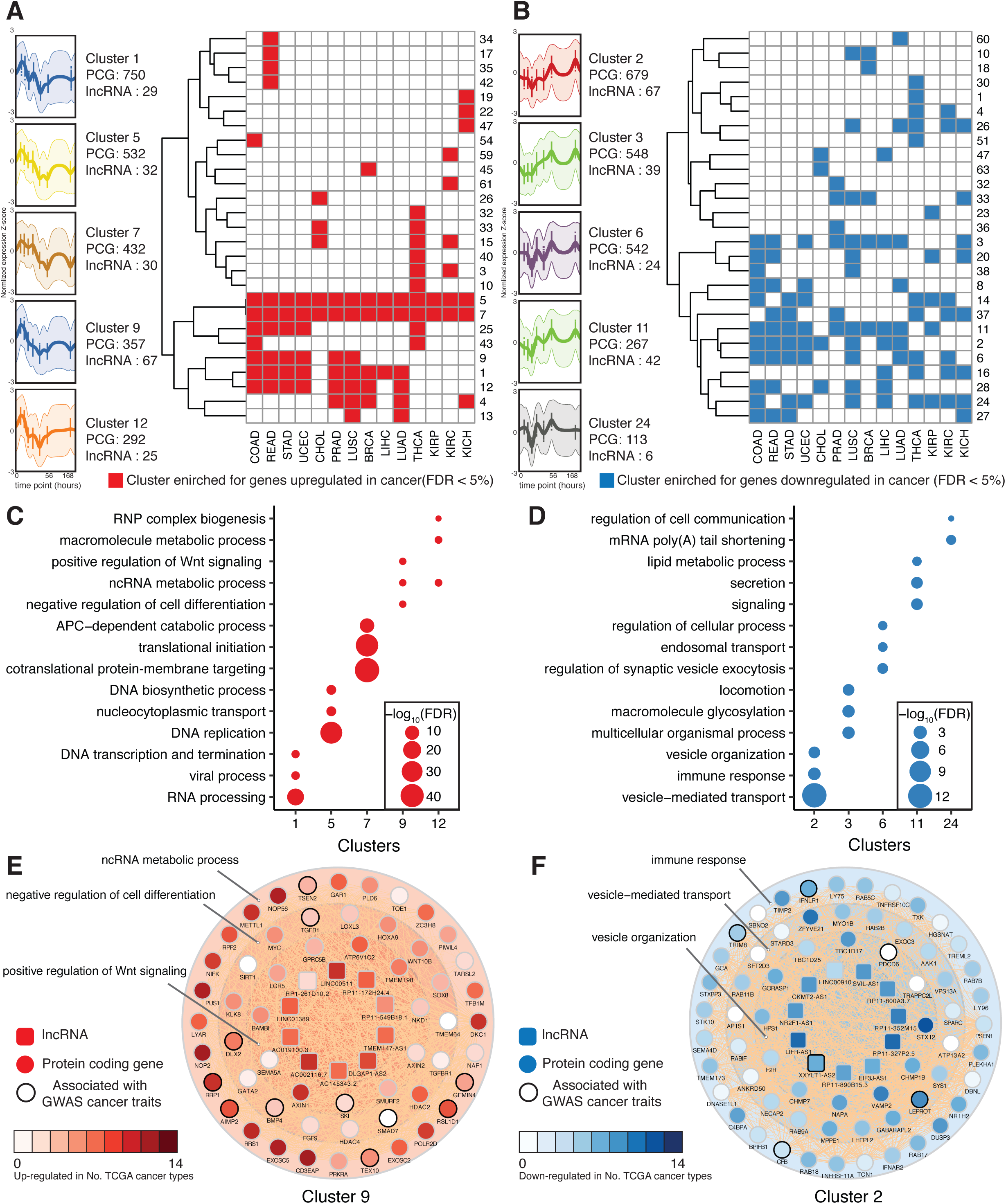
Wnt-regulated lncRNA and protein-coding genes form gene networks that are dysregulated in different cancer types. **(A)** Clusters enriched for genes upregulated in different cancer types. The top 5 clusters, cluster 1, 5, 7, 9 and 12 are enriched with the most number of cancers for genes upregulated. Normalized gene expression of these 5 clusters with number of PCGs and lncRNAs from each cluster are shown (left). **(B)** Clusters enriched for genes downregulated in different cancer types. The top 5 clusters, cluster 2, 3, 6, 11 and 24 are enriched with the most number of cancers for genes downregulated. Normalized gene expression of these 5 clusters with number of PCGs and lncRNAs from each cluster are shown (left). **(C, D)** GO Biological Processes enrichments (FDR < 5%) of the top 5 clusters enriched for genes upregulated (C) or downregulated (D) in different cancer types. The top 3 significantly enriched GO terms for each cluster are shown. (E) Wnt-regulated lncRNAs are part of gene networks that are upregulated in different cancers. PCGs from cluster 9 are enriched for ncRNA metabolic processes, negative regulation of cell differentiation and positive regulation of Wnt signaling. Wnt-regulated lncRNAs from cluster 9 are shown in the inner circle. (F) Wnt-regulated lncRNAs are part of gene networks that are downregulated in different cancers. PCGs from cluster 2 are enriched for immune response, vesicle-mediated transport and vesicle organization. Wnt-regulated lncRNAs from cluster 2 are shown in the inner circle.

### CRISPRi screens identify Wnt-regulated lncRNAs that modify HPAF-II cell growth in a context-dependent manner

Our analysis identified multiple Wnt-regulated lncRNAs, a subset of which might be important in cancer progression. To specifically identify the lncRNAs that play functional roles in the pathogenesis of *RNF43*-mutant pancreatic cancer *in vivo*, we performed CRISPRi screens. This approach utilizes dCas9-KRAB, where a catalytically inactive Cas9 is fused to a Krüppel associated box (KRAB) transcriptional repressor domain (Gilbert et al., 2013). dCas9-KRAB is recruited to the transcription start site (TSS) of lncRNAs by single guide RNAs (sgRNAs) to repress the transcription of the lncRNA of interest. CRISPRi screens have been demonstrated to be an efficient and specific approach for genome-wide loss-of-function studies of lncRNAs (S. J. Liu et al., 2017), which can not reliably be inactivated by indels introduced by the standard CRISPR-Cas9 system.

We chose to perform this CRISPRi screen *in vivo* because we have shown that both lncRNAs and PCGs respond to Wnt inhibition more robustly *in vivo* (Figure 1E-G and (Madan et al., 2018)), and that *in vivo* screening identifies dependencies not seen in tissue culture (Zhong et al., 2019). To capture the difference of Wnt-regulated lncRNA functions *in vivo* and *in vitro*, the CRISPRi screen was conducted both using xenograft tumor *in vivo* as well as cultured cells *in vitro* (Figure 4A).

**Figure 4:**
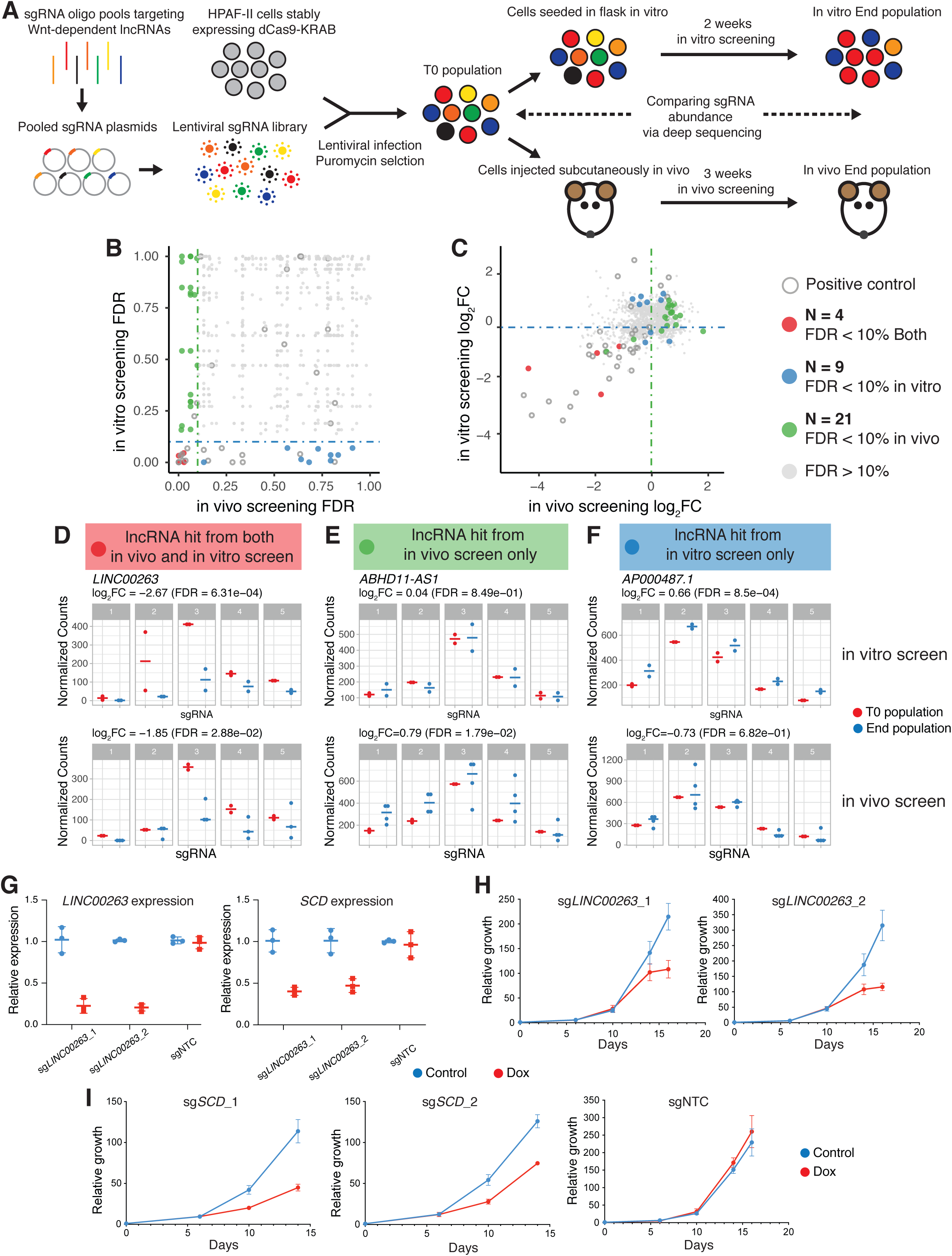
CRISPRi screens identify Wnt-regulated lncRNAs loci that modify cell growth in a context-dependent manner. **(A)** Schematic representation of CRISPRi screens conducted using xenograft tumors *in vivo* and in cultured cells *in vitro* to identify functional Wnt-regulated lncRNAs in *RNF43*-mutant pancreatic cancer. **(B)** Comparison of FDR from *in vivo* and *in vitro* screens. The dashed lines represent the threshold (FDR = 10%) for calling hits by gene-associated FDR. lncRNA hits are colored based on their FDR from both *in vivo* and *in vitro* screens. **(C).** Comparison of sgRNA fold change after *in vivo* and *in vitro* screens. Each gene is colored based on hits calling from B. **(D)** sgRNAs targeting *LINC00263* are significantly depleted from both *in vivo* and *in vitro* screens. **(E)** sgRNAs targeting *ABHD11-AS1* are significantly enriched only from the *in vivo* screen. **(F)** sgRNAs targeting *AP000487.1* are significantly enriched only from the *in vitro* screen. The normalized counts of 5 sgRNAs targeting the TSS of *LINC00263, ABHD11-AS1* and *AP000487.1* are shown before and after both screens in D, E, F. **(G)** sgRNAs targeting the TSS of *LINC00263* reduce the expression of *LINC00263 and SCD*. **(H)** sgRNAs targeting *LINC00263* reduce HPAF-II cell growth *in vitro*. Cell numbers were counted at days 6, 10, 14 and 16 after seeding and normalized to the seeding density. **(I)** sgRNAs targeting *SCD* reduce HPAF-II cell growth *in vitro*. sgNTC does not affect cell growth. Cell numbers were counted at day 6, 10 and 14 after seeding and normalized to the seeding density. NTC, non-targeting control.

We designed five sgRNAs to target the transcription start site (TSS) of each of the 1,503 Wnt-regulated lncRNAs (Horlbeck et al., 2016). We divided the sgRNAs into 3 lentiviral sub-libraries to allow for full representation of the sgRNAs throughout the *in vivo* screen, due to the limited number of cells that can be implanted and engrafted in each tumor. For each sub-library, we also included 55 sgRNAs targeting 11 genes involved in cell survival or Wnt signaling as positive controls, and 50 non-targeting controls (Table S4).

We transduced an HPAF-II cell line stably expressing dCas9-KRAB with the lentiviral sgRNA sub-libraries at a low multiplicity of infection (MOI < 0.3) to ensure that each cell was only infected by one virus with a single sgRNA. The transduced cells were selected with puromycin for 3 days (T0 population) and then maintained in culture for two weeks (the in vitro screen) with ≥ 3 x 10^6^ cells to allow for 1000-fold coverage of each sgRNA throughout the in vitro screen. Alternatively, the transduced cells were injected subcutaneously into immunocompromised mice. To get a good representation of each guide in the subcutaneous tumor, a total of 10^7^ cells were injected per mouse flank to allow for 3000-fold coverage of each sgRNA. The tumors were harvested after 3 weeks (the in vivo screen). Integrated lentiviruses encoding sgRNAs (i.e., barcodes) from the T0 population, the in vitro screen end population and the in vivo screen end population were then recovered by PCR and quantified by next-Gen sequencing (see Methods for additional details).

We first assessed the technical quality of the CRISPRi screen. There was a high correlation of sgRNA frequencies between independent experimental replicates (Figure S4), suggesting the robustness of the screen. We used the MAGeCK algorithm (W. Li et al., 2014) to analyze the *in vitro* and *in vivo* screens, using the non-targeting control sgRNAs for normalization. The statistical determination that a lncRNA gene regulated cancer proliferation was calculated based on the performance of all its sgRNAs compared to the non-targeting controls, as previously reported (W. Li et al., 2014). Each lncRNA gene was also scored based on the fold change of its second best performing sgRNA (W. Li et al., 2014). We classified a gene as a hit if its associated FDR was less than 10% (Figure 4B). First, our screen was able to identify important positive controls as gene hits. For example, all 5 sgRNAs targeting *POLR2A* (RNA polymerase II subunit A) were depleted in both *in vitro* and *in vivo* screens, consistent with its essential role for cell growth (Figure S5). As expected for a Wnt-addicted cancer, sgRNAs targeting *CTNNB1* were also depleted in both *in vitro* and *in vivo* screens (Figure S5). Thus, the screen appears to function well both *in vitro* and *in vivo*.

We next compared the lncRNA hits from the *in vivo* and *in vitro* screens. We identified 4 Wnt-regulated lncRNA loci as hits in both screens, 21 lncRNA loci as hits only in the *in vivo* screen and 9 lncRNA loci as hits only in the *in vitro* screen (Figure 4B and 4C, and Table 1). Since CRISPRi acts within a 1 kb window around the targeted TSS to repress gene expression (Gilbert et al., 2014), we also included in our sgRNA library guides designed to suppress the expression of the protein-coding genes that also had a TSS within 1 kb of the TSS of lncRNA hits. We found that for 6 lncRNA hits, protein-coding genes were nearby that could be suppressed by CRISPRi in the screen. However, CRISPRi suppression of these protein neighbors did not produce a phenotype in a separate screen library (Table S5). This indicates that the lncRNA hits identified through CRISPRi screen are likely due to the functions of lncRNA loci themselves. Taken together, around 1% (13/1,503) of the Wnt-regulated lncRNAs can modify cancer cell growth in the *in vitro* screen, which is consistent with previous genome-wide CRISPRi screens for functional lncRNAs in cell lines (S. J. Liu et al., 2017). Notably, using the *in vivo* CRISPRi screen, we identified twice as many Wnt-regulated lncRNAs (25/1,503) that had a functional effect on cancer cell growth.

**Table 1.**
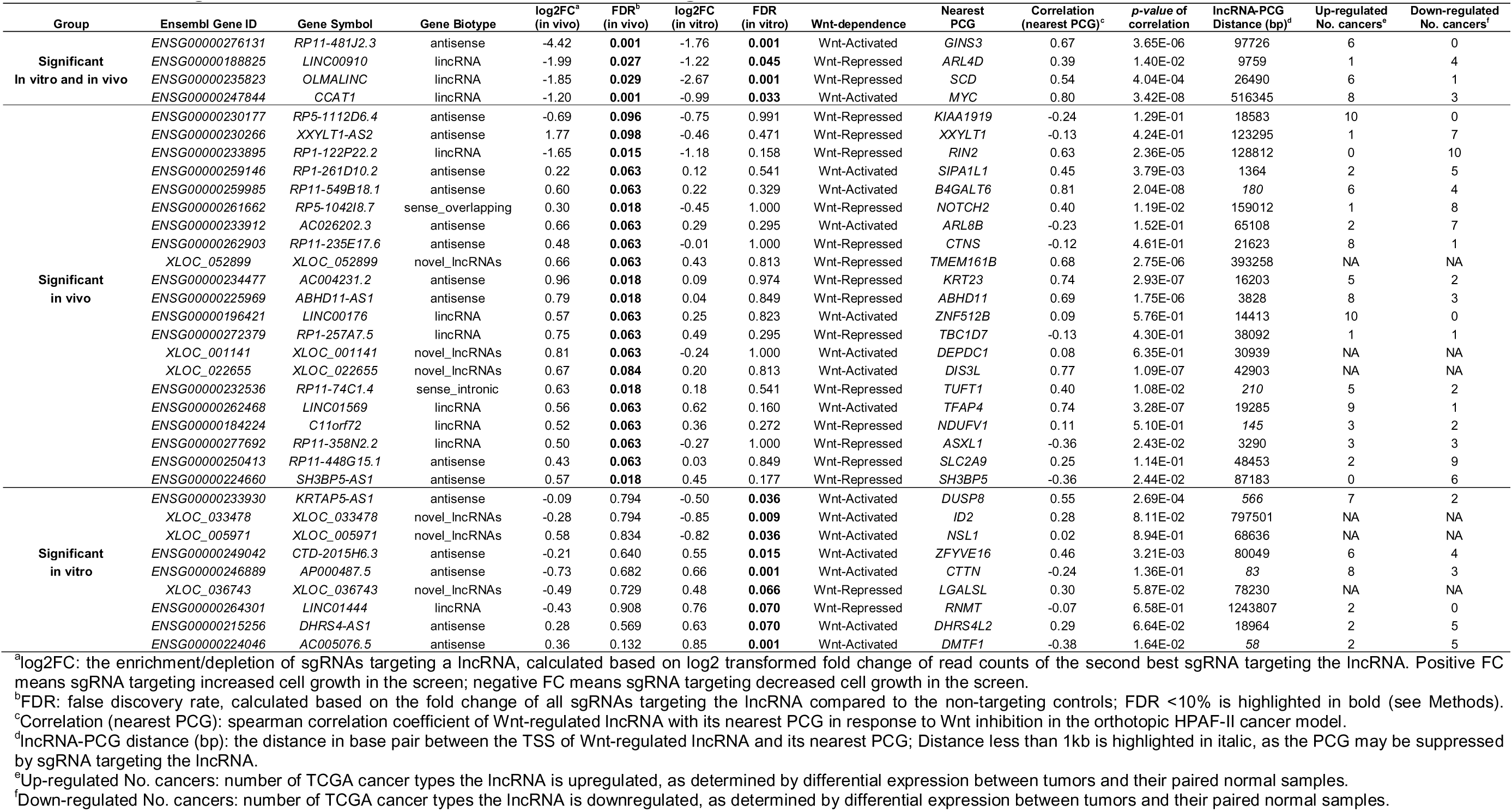
Wnt-regulated lncRNAs that affect HPAF-II cell growth in vivo and in vitro.

We found that the four Wnt-regulated lncRNA loci that were hits in both screens were essential for HPAF-II cancer cell growth (Figure 4B and 4C). For example, 3 out of 5 sgRNAs targeting *LINC00263* were depleted in both screens, suggesting that it was an essential lncRNA for HPAF-II growth both *in vivo* and *in vitro* (Figure 4D). Interestingly, *LINC00263* has previously been reported to be a cell type specific lncRNA essential for the growth of U87 cells but not K562, HeLa or MCF7 cells (S. J. Liu et al., 2017). 21 Wnt-regulated lncRNA loci were hits only in the *in vivo* screen and would not have been identified in an *in vitro* screen. Of these, 2 lncRNAs can promote cancer cell growth, while 19 lncRNAs appear to have suppressive effects on cell proliferation *in vivo*. For example, 4 sgRNAs targeting *ABGD11-AS1* were only enriched at the end of the *in vivo*, but not the *in vitro* screen (Figure 4E). Among the 9 Wnt-regulated lncRNA loci that were hits only in the *in vitro* screen, we found 3 of them promoted, while 6 suppressed HPAF-II proliferation in culture. For example, all 5 sgRNAs targeting *AP000487.1* were enriched at the end of the *in vitro* screen, however, none of the 5 sgRNAs showed significant change after the *in vivo* screen (Figure 4F). This suggests that *AP000487.1* may have tumor suppressive function only *in vitro*. Taken together, using CRISPRi screens both *in vivo* and *in vitro*, we identified Wnt-regulated lncRNAs loci that modify HPAF-II growth in a context-dependent protein-coding genes that also had a TSS within 1 kb of the TSS of lncRNA hits manner. It also suggests that lncRNA loci identified *in vitro* may not have important functions *in vivo*.

To further validate the CRISPRi screen results, we focused on *LINC00263*, which was an essential lncRNA for HPAF-II cell growth both *in vivo* and *in vitro* (Figure 4D). We cloned the top two sgRNAs targeting *LINC00263* into doxycycline-inducible lentiviral sgRNA vectors. After confirming the doxycycline-inducible knockdown of *LINC00263* expression in the HPAF-II cell lines (Figure 4G), we verified that knocking down *LINC00263* reduced HPAF-II cell growth *in vitro* (Figure 4H). Interestingly, we found that knocking down *LINC00263* also reduced the expression of its nearest protein-coding gene stearoyl-CoA desaturase (*SCD*) (Figure 4G), similar to what was reported in U87 cells (S. J. Liu et al., 2017). To test if *SCD* regulates the growth of HPAF-II cells, we next targeted the TSS of *SCD* using CRISPRi with two independent sgRNAs. Knockdown of *SCD* reduced *SCD* mRNA abundance (Figure S6) and inhibited HPAF-II cell growth similar to that observed after knockdown of *LINC00263* (Figure 4I). However, sgRNAs targeting the TSS of *SCD* did not reduce the expression of *LINC00263* (Figure S6). Based on these results, we hypothesize that *LINC00263* is essential for HPAF-II cell growth through *cis*-regulation of *SCD*.

## Discussion

LncRNAs play important roles in diverse biological processes. Here we present a systematic study to identify and functionally assess lncRNAs regulated by Wnt signaling. Using an orthotopic Wnt-addicted pancreatic cancer model treated with a potent and effective PORCN inhibitor, we identified 1,503 lncRNAs regulated by Wnt signaling *in vivo*. Many of these lncRNAs were also dysregulated in different cancer types and may function in gene networks that contribute to the pathogenesis of cancers. Our eQTL-lncRNA interactions analysis identified Wnt-regulated lncRNAs that may regulate nearby protein-coding genes. Using CRISPRi screens, we found that 34 Wnt-regulated lncRNAs could modify cell growth in a context-dependent manner with a higher hit rate in the *in vivo* model. This pipeline for lncRNA discovery and functional validation may be broadly applicable.

We previously reported that Wnt-regulated protein-coding genes were more robustly regulated in an orthotopic model than in cultured cells. We find that this holds true for lncRNAs as well. More than twice as many lncRNAs responded to Wnt inhibition in the *in vivo* xenografts than in cells cultured *in vitro*. These differences in the number and magnitude of gene expression changes will be influenced by a variety of local and experimental factors including tumor microenvironment, culture conditions, doubling times in different environments, local nutrients versus culture medium ingredients, the presence of stromal and other host cells, and variations in extracellular matrix. Overall, our findings are consistent with the large body of literature showing that the expression of genes is regulated by interaction with the relevant environment (Killion et al., 1998).

Cancer cells show differential dependencies on protein-coding genes for their growth and survival *in vivo* versus *in vitro* (Miller et al., 2017; Possik et al., 2014; Yau et al., 2017; Zhong et al., 2019). Our CRISPRi screen results indicate that cancer cells also have different requirements for lncRNAs when grown *in vivo* vs *in vitro* conditions. Multiple lncRNAs exhibit different phenotypes when studied in cell culture compared to animal knock-out models and *in vivo* systems (Bassett et al., 2014; Goudarzi et al., 2019; Han et al., 2018; Kohtz, 2014; Ruan et al., 2020). Our results highlight the importance of studying lncRNAs *in vivo* with the relevant microenvironment in order to better understand their functions in cancer pathogenesis. This has implications for the identification of lncRNAs as potential therapeutic targets for cancer treatment. For instance, it has been shown that drugs identified through high-throughput screening of cell culture *in vitro* have limited success in patient care (Letai, 2017; Sharma et al., 2010). The same might be true for drugs identified to target lncRNAs.

Despite the large number of lncRNAs annotated in the human genome (Hon et al., 2017; Iyer et al., 2015), only a very small fraction of them have been either validated or characterized at a functional level. This is due to the complex nature of the lncRNA loci and a prior lack of tools to study them at a large scale (Bassett et al., 2014; Kopp & Mendell, 2018). In recent years, CRISPR screens have been shown to be an efficient and specific approach to investigate lncRNA functions genome-wide in cultured cells (Esposito et al., 2019; Joung et al., 2017; S. J. Liu et al., 2017; Zhu et al., 2016). In this study, we perform a CRISPRi screen not only in cultured cells, but also in xenograft tumors to assess the ability of 1,503 Wnt-regulated lncRNAs to influence cancer cell proliferation. Validating this approach, among the 4 Wnt-regulated lncRNAs that we found to be functional both *in vivo* and *in vitro*, 3 were identified to promote cell growth in prior CRISPRi screens (S. J. Liu et al., 2017). Furthermore, consistent with what has been reported for genome-wide lncRNA CRISPRi screens in cell lines (S. J. Liu et al., 2017) 1% (13/1,503) of the Wnt-regulated lncRNAs in our *in vitro* screen modified cancer cell growth. Notably, our *in vivo* CRISPRi screen identified twice as many Wnt-regulated lncRNAs (25/1,503) that had a functional effect on cancer cell growth. 21 Wnt-regulated lncRNAs had functional effects on cancer cell growth only in the xenograft model and would not have been identified in an *in vitro* screen, demonstrating the value of studying lncRNA functions *in vivo*. This is also demonstrated in a recent study that an *in vivo* system is essential for understanding the biological role of a human lncRNA in metabolic regulation that cannot be recapitulated *in vitro* (Ruan et al., 2020).

The CRISPR based approach can produce different results than those based on RNA interference. *LINC00176*, found in our screen as a functional Wnt-regulated lncRNA locus, has also been identified in four other publications. Two groups used different CRISPR approaches (paired-sgRNAs (Zhu et al., 2016) or sgRNA targeting splice site (Y. Liu et al., 2018)) and found, as we did, that *LINC00176* has a tumor-suppressive effect *in vivo*. Two additional studies used RNA interference and concluded, conversely, that *LINC00176* has a pro-proliferative role in ovarian and hepatocellular carcinoma cell lines (Dai et al., 2020; Tran et al., 2018). These differences could be due to differences in cell type or experimental approach, as RNA interference is known to suffer from significantly more off-target effects compared to the CRISPR approach and is less effective for targeting nuclear lncRNAs (Smith et al., 2017; Stojic et al., 2018). Together, the comparisons here further support the identification of Wnt-regulated lncRNA loci that can modify cancer cell growth and the importance of choosing a loss-of-function strategy to characterize lncRNAs.

Nevertheless, there are some limitations to using CRISPRi to target lncRNAs. First, recruiting dCas9-KRAB to the TSS of a lncRNA can suppress the transcriptional activity and local regulatory sequence (enhancer) of the lncRNA locus; second, it results in decreased production of the lncRNA transcript, inhibiting potential *cis* or *trans* function of the lncRNA transcript (S. J. Liu et al., 2017). Both the repressive effect on chromatin and the lack of lncRNA transcripts can cause biological consequences that cannot be differentiated by CRISPRi knock-down alone. Thus, additional studies are needed to dissect how the Wnt-regulated lncRNA loci identified in our screen regulate cell proliferation.

GWAS studies have identified thousands of common genetic variants that are associated with complex traits and diseases, but 90% of these fall into noncoding regions of the genome (Hindorff et al., 2009). This has made it difficult to dissect the underlying molecular mechanisms. eQTLs that co-localize with GWAS SNPs suggest the effect of the SNPs on diseases and traits is mediated by changes in gene expression. lncRNAs overlapping with these GWAS associated *cis*-eQTL SNPs are potential candidates to explain the underlying mechanisms of risk loci because lncRNAs can be important *cis* regulators of nearby genes (Engreitz et al., 2016; Gil & Ulitsky, 2019; Luo et al., 2016). When we mapped Wnt-regulated lncRNAs-mRNA pairs linked by eQTL SNPs using the annotation from FANTOM5 (Hon et al., 2017) we found previously unappreciated regulatory effects of Wnt-regulated lncRNAs in disease. For example, Wnt-regulated lncRNA *LINC00339* was linked to *CDC42* through five eQTL SNPs, suggesting the *LINC00339* locus may regulate the expression of *CDC42*. Supporting this, knocking-down *LINC00339* expression has been reported to increase *CDC42* expression (X.-F. Chen et al., 2018). Consistent with the importance of Wnt regulation, *LINC00339* and its linked gene *CDC42* are involved in both endometriosis and bone metabolism (X.-F. Chen et al., 2018; Powell et al., 2016), two Wnt-regulated biological processes (Krishnan, 2006; Yongyi Wang et al., 2009). Thus, identifying eQTL-linked Wnt-regulated lncRNA-PCG pairs helps to prioritize the potential *cis*-regulatory targets of Wnt-regulated lncRNAs. Further integrating the disease risk information based on GWAS SNPs co-localizing with eQTL, the Wnt-regulated lncRNA-PCG pairs may help explain the underlying mechanisms of risk loci in the context of disease, which is potentially affected by Wnt signaling.

Although the 1,503 Wnt-regulated lncRNAs were discovered in the orthotopic *RNF43-*mutant pancreatic cancer xenograft model, many of them were also dysregulated in different types of cancers in TCGA (Figure S1A). 253 Wnt-regulated lncRNAs were exclusively upregulated or downregulated across different cancer types (Table S1). This suggests the fundamental roles of Wnt-regulated lncRNAs in cancer pathogenesis in a broader context beyond Wnt-addicted pancreatic cancer. For example, *CCAT1,* identified as a Wnt-activated lncRNA, was also upregulated in 9 cancer types (Table S1). Our CRISPRi screens indicated that it is an essential lncRNA both *in vivo* and *in vitro* (Table 1). This suggests that *CCAT1* is a Wnt-activated lncRNA with oncogenic function, which is consistent with previous studies showing that *CCAT1* can promote the progression of different types of cancers (Y. Jiang et al., 2018; Xiang et al., 2014; E. Zhang et al., 2017). Integrating Wnt-regulated lncRNAs with their expression profiles in TCGA and CRISPRi functional screens can better distinguish their oncogenic or tumor suppressive functions in cancer pathogenesis.

## Conclusions

This study comprehensively identified 1,503 lncRNAs regulated by Wnt signaling *in vivo* and determined their wider roles in other cancers. We found more than twice as many lncRNAs responded to Wnt inhibition in the *in vivo* xenografts than in cells cultured *in vitro*. With CRISPRi screens both *in vivo* and *in vitro*, we found two fold (21/1503) as many Wnt-regulated lncRNAs have functional effects on cell growth only *in vivo*, suggesting the importance of studying lncRNA function with relevant microenvironment. Thus, this study provides a valuable resource of functional Wnt-regulated lncRNAs *in vivo*. It also establishes a framework for integrating orthogonal transcriptomics dataset with functional CRISPRi screening which can be broadly adapted for systematic discovery, functional annotation and validation of lncRNAs *in vivo*.

## Methods

### De novo lncRNA discovery

The polyA+ RNA-seq dataset contains the transcriptional response to PORCN inhibitor ETC-159 treatment at seven time points (0, 3, 8, 16, 32, 56 and 168 hours) using an orthotopic model of *RNF43*-mutant pancreatic adenocarcinoma (HPAF-II). The data was previously published (Madan et al., 2018) under accession number GSE118041. RNA-seq reads were assessed for quality with FASTQC. Reads originating from mouse genome (mm10) were removed with Xenome (Conway et al., 2012). All the reads among replicates from each time point were pooled to achieve deep coverage for novel lncRNA discovery. Each time point generated between 160 million to 237 million reads. The reads were aligned to hg38 (Ensembl version 79) using TopHat v2.0.10 (Kim et al., 2013). De novo transcriptome assembly was performed separately for each time point with Cufflinks v2.1.1 (Trapnell et al., 2010). Transcriptome assemblies at each time point were merged and compared with Ensembl build 79 as reference, using Cuffmerge. The novel transcripts were selected using Cuffcompare class code for novel intergenic and novel antisense transcripts. All the novel transcripts were then merged with Ensembl build 79 to establish a full reference transcriptome. RNA-seq reads from each sample were also individually aligned to hg38 (Ensembl version 79) using TopHat v2.0.10 (Kim et al., 2013). Gene level reads counts for each sample were computed with HTSeq 0.6.0 (Anders et al., 2015), which were then converted to gene expression in Transcripts per Million (TPM). To identify putative novel lncRNAs transcripts, the novel transcripts were filtered using the following criteria: length longer than 200 bp and estimation to be non-protein coding based on three methods: CPAT with threshold less than 0.364 (L. Wang et al., 2013), CPC with threshold less than 0 (Kong et al., 2007) and Slncky defined as “lncRNA” (J. Chen et al., 2016). Known lncRNAs from Ensembl build 79 were obtained based on their transcript biotype: “lincRNA”, “antisense”, “sense_intronic”, “sense_overlapping”. All the genes were also filtered based on their expression to make sure that the median expression level of each gene at every time point had TPM > 1. This analysis yielded 16,160 genes, including 12,527 protein-coding genes, 2,846 annotated lncRNAs and 787 novel lncRNAs that were expressed in *RNF43*-mutant pancreatic adenocarcinoma (HPAF-II).

### Identification of Wnt-regulated lncRNAs

To identify genes regulated by Wnt signaling, DESeq2 (Love et al., 2014) was used to perform differential expression analysis on 16,160 genes across time points with likelihood ratio test (LRT). Adjusted *P* value < 0.05 was used to select genes significantly responded to Wnt inhibition across time points. This led to 10,554 Wnt-regulated genes, including 9,051 protein-coding genes and 1,503 lncRNAs (1,178 annotated lncRNAs and 325 novel lncRNAs).

### Comparison of lncRNAs response to Wnt inhibition across models

Two RNA-seq datasets contain transcriptional response of *in vitro* model (48 h ETC and 48 h Veh) and subcutaneous model (0h and 56h) of *RNF43*-mutant pancreatic adenocarcinoma (HPAF-II) to PORCN inhibitor ETC-159 treatment. The data was previously published (Madan et al., 2018) under accession number GSE118190 and GSE118179, respectively. RNA-seq reads from these datasets were assessed for quality with FASTQC (https://www.bioinformatics.babraham.ac.uk/projects/fastqc/). Reads originating from mouse genome (mm10) were removed with Xenome (Conway et al., 2012) and aligned to hg38 (Ensembl version 79) using TopHat v2.0.10 (Kim et al., 2013) for each sample. Gene level reads counts were computed with HTSeq 0.5 (Anders et al., 2015). DESeq2 (Love et al., 2014) was used to perform differential gene expression analysis on 16,160 genes between the time points with Wald test for each of the models, namely *in vitro* model (48 h ETC and 48 h Veh), subcutaneous model (0h and 56h) and orthotopic model (0h and 56h). An adjusted *P* value < 0.1 was used to select genes that significantly responded to Wnt inhibition between the two time points.

### Wnt-regulated lncRNA co-expression with PCGs

The degree of co-expression between Wnt-regulated lncRNAs and either all PCGs or their nearest PCG in response to Wnt inhibition in the orthotopic HPAF-II cancer model was calculated with cor function (spearman correlation) in R. The TAD data from the PANC-1 cell line mapped to hg38 was downloaded from the 3D Genome Browser (Yanli Wang et al., 2018). The Wnt-regulated lncRNA and nearest PCG pair were classified into two groups, the pair in the same TAD versus the pair in different TADs based on the PANC-1 TAD information. The correlation distributions between the two groups were tested for difference by using a two-sample nonparametric Mann–Whitney U test using the R function wilcox.test.

### Analysis of TCGA dataset

HTSeq - Counts data of all the TCGA cancers were downloaded from the UCSC Xena platform (Goldman et al., 2019). The cancer types were selected for further analysis if at least 5 tumor-normal pairs were present, and there was a clear separation between the tumor and normal samples in the dataset based on PCA analysis. This yielded 14 cancer types. Genes with less than 10 reads mapped across the samples within each cancer type were removed. Differential expression analysis between the paired tumor-normal samples for each cancer type was performed using DESeq2 (Love et al., 2014). An adjusted *P* value < 0.05 was used to select genes significantly differentially expressed between tumor and normal sample.

### Integrative analysis of FANTOM5 dataset

Wnt-regulated lncRNAs were mapped to FANTOM5 lncRNA annotations as follows:

1. If the lncRNA was annotated with the same Ensembl Gene ID in FANTOM5, it’s considered the same lncRNA. 2. The remaining lncRNAs were overlapped with FANTOM5 lncRNA assembly (hg38) to identify the corresponding FANTOM5 CAT_geneID. Among the 1,503 Wnt-regulated lncRNAs, 1,073 were also annotated in FANTOM5 and 430 were novel previously unannotated lncRNAs. The eQTL linked lncRNA protein-coding gene (PCG) pairs for these 1,073 annotated Wnt-regulated lncRNAs were extracted from FANTOM5 annotation eQTL_linked_lncRNA_mRNA_pair (Hon et al., 2017). This yielded 1,486 lncRNA-PCG mRNA pairs linked by eQTL SNPs involving 602 Wnt-regulated lncRNAs (Figure 2A and Table S2). The gene expression profiles of all the pairs in 1,829 FANTOM5 samples were downloaded from the expression atlas FANTOM_CAT.expression_atlas.gene.lv3_robust.rle_cpm curated by FANTOM5 (Hon et al., 2017). The lncRNA-PCG pair was identified as significantly co-expressed in FANTOM5 samples if it passed the threshold used in (Hon et al., 2017), i.e., that their co-expression is greater than 75th percentile of the matched background correlation (*binom_p* < 0.05 compared to the background). The lncRNA-PCG pair co-expression in response to Wnt inhibition was calculated using Spearman correlation *rho* on gene expression TPM across time points. The associated *p* value was also calculated using cor.test function in R. To identify the eQTL that are co-localizing with GWAS SNP, eQTLs linking Wnt-regulated lncRNA and protein-coding genes were first mapped to SNP id using biomart in R. These SNPs were overlapped with trait-associated SNPs curated by FANTOM5 to subest the SNPs associated with cancer by GWAS. In total, 271 eQTL SNPs were found to be associated with cancer by GWAS, linking 115 Wnt-regulated lncRNA-PCG pairs involving 49 Wnt-regulated lncRNAs (Table S3).

### Time series clustering

Time series clustering on 10,554 Wnt-regulated genes was performed using GPClust (Hensman et al., 2013) as previously described (Madan et al., 2018). Gene expression TPM were converted to z-scores and time points were square root transformed. Genes were clustered with GPClust (Hensman et al., 2013) using the Matern32 kernel with a length scale of 6 and a concentration (alpha) parameter of 0.001, 0.01, 0.1, 1, and 10. Genes were assigned to a cluster based on the highest probability of being a member of that cluster. Clustering was performed 10 times for a specified set of parameters, with the best clustering taken as the one with the lowest distance to the other clusterings, i.e. the most representative.

### Functional enrichment analysis

Gene Ontology (GO) enrichment analysis was performed with g:Profiler (Reimand et al., 2016) using all the Wnt-regulated protein-coding genes as background. Significantly enriched GO terms were selected with FDR < 5%.

### Enrichment analysis for dysregulated genes from different cancers

Genes significantly differentially expressed (adjusted *P* value < 0.05) between tumor-normal pairs were defined as dysregulated genes. To test whether the clusters were enriched for dysregulated genes in each cancer type, genes from each of the 63 clusters were intersected with dysregulated genes from each cancer separately by carrying out a Fisher’s exact test. The gene background used for the test were Wnt-regulated genes that were dysregulated in the specific cancer. Upregulated genes and downregulated genes were examined separately for enrichment. The Fisher’s exact test was performed with fisher.test in R for overrepresentation. Nominal *p* values were adjusted for multiple testing using the Benjamini-Hochberg method. Clusters significantly enriched for dysregulated genes were selected with FDR < 5%. The significance of the enrichment was clustered for each cluster and its enriched cancer type.

### CRISPRi sgRNA library design

CRISPRi single guide RNA (sgRNA) library was designed to target the transcription start site (TSS) of each of the Wnt-regulated lncRNAs. 1,503 Wnt-regulated lncRNAs were selected for the CRISPRi screen, which contained 3,151 transcripts including different isoforms. To avoid redundancy of different TSSs located in close proximity, if TSSs of transcripts belonging to the same gene were within 100bp, they were grouped together. A total set of 2,337 TSSs were obtained for Wnt-regulated lncRNAs, which were then converted to hg19 with the liftover function in R. These TSSs were furthered refined with FANTOM based TSS annotation and 5 sgRNAs were designed to target each of the TSS using hCRISPRi-v2.1 algorithm (Horlbeck et al., 2016). Since some TSSs could not be uniquely targeted, in total 8,560 sgRNAs were designed to target 1,486 Wnt-regulated lncRNAs. The sgRNAs were then divided into 3 sub-libraries. Protein-coding genes whose TSSs were within 10 kb of Wnt-regulated lncRNAs were selected. sgRNAs targeting these protein-coding genes were extracted from the hCRISPRiv2 library (Horlbeck et al., 2016) to constitute a 4th sub-library. For each sub-library we also included 55 sgRNAs targeting 11 genes (*PCNA*, POLR2A, *PSMA7*, *RPS27, SF3A3, CTNNB1, FZD5, APC, AXIN1, CSNK1A1, PORCN*) involved in cell survival and Wnt signaling as positive controls and 50 non-targeting controls (Table S4). The sgRNAs libraries were synthesized by CustomArray (Bothell, WA, USA).

### sgRNA cloning and lentiviral packaging

The sgRNA libraries were cloned into pCRISPRia-v2 sgRNA expression vector (Horlbeck et al., 2016) by Gibson assembly (NEB). They were then amplified using electroporation in Endura electrocompetent cells (Lucigen), to achieve at least 250 colonies per sgRNA in the library. For individual CRISPRi knockdown, top 2 performing sgRNAs targeting *LINC00263* and *SCD* were selected with protospacer sequences: sg*LINC00263_1* (GACCTCAGTCTGCCCTACCC), sg*LINC00263_2* (GGGTAGGGCAGACTGAGGTC), sg*SCD*_1 (GCTTGGCAGCGGATAAAAGG), sg*SCD*_2 (GCACATTCCCAACTCACGGA). The sgRNAs were cloned into doxycycline-inducible lentiviral sgRNA expression vector FgH1tUTG as previously described (Aubrey et al., 2015). The sgRNA plasmid was packaged into lentiviral particles with psPAX2 and pMD2.G packaging plasmids. The virus supernatant was harvested 48 and 72 hours after transfection, filtered through 0.45 µm filter and stored at -80 °C.

### Cell lines

The HPAF-II cell line was obtained from the Duke Cell Culture Facility. An HPAF-II stable cell line expressing dCas9-KRAB was generated by lentiviral transduction with pMH0001 plasmid (UCOE-SFFV-dCas9-BFP-KRAB) (Adamson et al., 2016) and sorting for the top 20% −30% BFP expressing cells. All cell lines were cultured in Eagle’s Minimum Essential Medium (EMEM) supplemented with 10% FBS, 1 mM sodium pyruvate, 2 mM L-glutamine and 10% penicillin/streptomycin, maintained in 5% CO_2_. Cells were regularly tested for mycoplasma.

### CRISPRi screens

The HPAF-II-dCas9-KRAB stable cell line was infected with sgRNA lentiviral libraries at a multiplicity of infection (MOI) < 0.3 with 8 µg/ml polybrene. The infected cells were selected with 2 µg/ml puromycin for 3 days (T0 population). 3 x 10^6^ cells from the T0 population were harvested and stored as a cell pellet at −20 °C for sequencing. For the *in vitro* screen, cells from T0 population were passaged with a seeding density of 3 x 10^6^ cells at each passage to allow for 1000 times coverage of each sgRNA, and cultured for 2 weeks. 3 x 10^6^ cells at the end of the *in vitro* screen were harvested and stored as a cell pellet at −20 °C for sequencing. The *in vitro* screen was performed in duplicates for each sub-library. For the *in vivo* screen, cells from the T0 population were mixed with Matrigel (BD Biosciences) and injected subcutaneously into the flanks of NOD-scid gamma (NSG) mice. 10^7^ cells were injected per flank to allow for library coverage of 3000 cells/sgRNA at the time of implantation. A group of 3 mice were injected per sub-library. Mice were sacrificed 3 weeks after injection, and tumors were harvested and stored at −80°C. Genomic DNA from the frozen cell pellets and homogenized tumors was extracted with high salt precipitation. The sgRNA region was amplified by PCR. A second round of PCR was performed to append Illumina sequencing adaptors and barcodes for each sample. PCR products were purified and quantified with a Bioanalyzer, and sequenced on the Illumina MiSeq platform.

### CRISPRi screens analysis

Reads from sequenced screening sgRNA libraries were demultiplexed based on sample barcodes with FASTX-Toolkit. The reads were then counted against individual sub-libraries using MAGeCK count function (W. Li et al., 2014) with non-targeting control sgRNA for normalization. sgRNA counts were used for quality control using PCA and clustering analysis with DESeq2 (Love et al., 2014) to exclude outlier samples. Robust Rank Aggregation analysis (RRA) was performed with MAGeCK (W. Li et al., 2014) test function to detect sgRNAs significantly depleted or enriched from the screens. Gene level significance was calculated based on the performance of all its sgRNAs compared to non-targeting controls, as previously shown (W. Li et al., 2014). Each gene was also scored based on the fold change of its second best performing sgRNA <u>(W. Li et al.,2014)</u>. We classified genes as hits if their associated FDR < 10%.

### Inducible CRISPRi knockdown

1 µg/ml doxycycline final concentration (dox) (from a stock of 10 mg/ml dissolved in DMSO) was used to induce sgRNA expression from the inducible lentiviral sgRNA expression vector, while DMSO was used as the control. After 48 hours induction, total RNA was isolated from the CRISPRi knockdown cells. RT-qPCR was performed to assess the knockdown efficiency for *LINC00263 and SCD with HPRT* gene as an internal control. RT-qPCR primers were: *LINC00263_*Forward (AAAGATTGGGCAGTCACTGG)*, LINC00263_*Reverse (TGGGTCTTCAGCACCAAATG)*, SCD_*Forward (TTCCTACCTGCAAGTTCTACACC)*, SCD_*Reverse (CCGAGCTTTGTAAGAGCGGT). The effect of CRISPRi knockdown on cell growth was assessed with internally controlled, relative growth assays. Cells were seeded in duplicates and treated with either 1 µg/ml dox or DMSO. Cells were counted every 3-4 days after the initial dox treatment.

## Additional Files

**Additional file 1: Table 1.** Wnt-regulated lncRNAs that affect HPAF-II cell growth in vivo and in vitro. (DOCX 26 kb)

**Additional file 2: Figure S1.** Wnt-regulated lncRNAs and PCGs are dysregulated in TCGA cancers (A) Wnt-regulated lncRNAs and PCGs, defined as genes changed over time upon Wnt inhibition (FDR < 5%) in the orthotopic RNF43-mutant pancreatic cancer model (Figure 1A). Wnt-regulated lncRNAs and PCGs are dysregulated in different types of cancers as determined by differential expression between tumors and their paired normal samples using the TCGA dataset. (B) VPS9D-AS1 is upregulated in 11 different types of cancers. (PDF 946 kb)

**Additional file 3: Figure S2.** Subset of Wnt-dependent lncRNAs co-express with its nearest PCG in the same TAD. (A) Wnt-regulated lncRNAs exhibit stronger co-expression with their nearest PCG after Wnt inhibition compared to their co-expression with all PCGs. (B) Wnt-regulated lncRNA–nearest PCG pairs within the same TAD exhibit stronger co-expression than the pairs in different TADs. P for significance was calculated by Mann–Whitney U test. (C) For the Wnt-regulated lncRNA–nearest PCG pairs encoded within the same TAD, the PCGs are significantly (FDR < 5%) enriched for GO biological processes. (1076 kb)

**Additional file 4: Figure S3.** Clusters are enriched for genes dysregulated in different cancers. (A) The Wnt-regulated lncRNAs and PCGs fall into 63 distinct clusters based on their pattern of expression change following Wnt inhibition. (B) 46 out of the 63 clusters are enriched (FDR < 5%) for genes dysregulated in at least one type of cancer. (PDF 5287 kb)

**Additional file 5: Figure S4.** A high correlation of sgRNA counts between independent experimental replicates in CRISPRi screens. (A) Correlation of sgRNA counts between experimental replicates in the in vitro screens. (B) Correlation of sgRNA counts between experimental replicates in the in vivo screens. (PDF 959 kb)

**Additional file 6: Figure S5.** CRISPRi screens are able to identify important positive controls as gene hits. (PDF 909 kb)

**Additional file 7: Figure S6.** Knockdown of SCD with CRISPRi reduce SCD mRNA abundance, but not the expression of LINC00263. (PDF 876 kb)

**Additional file 8: Table S1.** 1,503 Wnt-regulated lncRNAs. (XLSX 139 kb)

**Additional file 9: Table S2.** Wnt-regulated lncRNA-PCG pairs linked by eQTL SNPs involving 602 Wnt-regulated lncRNAs. (XLSX 158 kb)

**Additional file 10: Table S3.** Wnt-regulated lncRNAs were linked by eQTL SNPs that colocalize with cancer GWAS loci. (XLSX 63 kb)

**Additional file 11: Table S4.** sgRNA libraries used in CRISPRi screens. (XLSX 767 kb)

**Additional file 12: Table S5.** CRISPRi screens results on protein-coding gens that have TSS within 1 kb of the TSS of Wnt-dependent lncRNAs. (XLSX 135 kb)

## Declarations

Ethics approval and consent to participate

This work has been approved by the Duke-NUS Institutional Animal Care and Use Committee.

## Consent for publication

Not applicable

## Availability of data and materials

The RNA-seq data for HPAF-II orthotopic, subcutaneous and *in vitro* model is available at NCBI GSE118041, GSE118179, GSE118190. Detailed results for the Wnt-regulated lncRNAs can be found in supplementary tables.

## Competing interests

BM and DMV have a financial interest in ETC-159.

## Funding

This study is supported in part by the National Research Foundation Singapore and administered by the Singapore Ministry of Health’s National Medical Research Council under the STAR Award Program to DMV. EP acknowledges the support of Duke-NUS Medical School, Singapore. BM acknowledges the support of the Singapore Ministry of Health’s National Medical Research Council Open Fund–Independent Research Grant.

## Authors’ contributions

SYL, DMV, EP conceived the project. SYL performed the data analysis with assistance from NH. SYL performed the CRISPRi screens with assistance from YKW, BM and ZZ. SYL and TLG performed the inducible CRISPRi knockdown validation with assistance from ZZ. SYL, DMV, EP wrote the manuscript with inputs from NH and BM. All authors read and approved the final manuscript.

## Acknowledgements

We thank the members of the DMV and EP labs and the Duke-NUS scientific community for their helpful comments and discussion

## References

1. Adamson, B., Norman, T. M., Jost, M., Cho, M. Y., Nuñez, J. K., Chen, Y., Villalta, J. E., Gilbert, A., Horlbeck, M. A., Hein, M. Y., Pak, R. A., Gray, A. N., Gross, C. A., Dixit, A., Parnas, O., Regev, A., & Weissman, J. S. (2016). A Multiplexed Single-Cell CRISPR Screening Platform Enables Systematic Dissection of the Unfolded Protein Response. Cell, 167(7), 1867–1882.e21.

2. Anders, S., Pyl, P. T., & Huber, W. (2015). HTSeq--a Python framework to work with high-throughput sequencing data. Bioinformatics, 31(2), 166–169.

3. Aubrey, B. J., Kelly, G. L., Kueh, A. J., Brennan, M. S., O’Connor, L., Milla, L., Wilcox, S., Tai, L., Strasser, A., & Herold, M. J. (2015). An inducible lentiviral guide RNA platform enables the identification of tumor-essential genes and tumor-promoting mutations in vivo. Cell Reports, 10(8), 1422–1432.

4. Bailey, P., Chang, D. K., Nones, K., Johns, A. L., Patch, A.-M., Gingras, M.-C., Miller, D. K., Christ, A. N., Bruxner, T. J. C., Quinn, M. C., Nourse, C., Murtaugh, L. C., Harliwong, I., Idrisoglu, S., Manning, S., Nourbakhsh, E., Wani, S., Fink, L., Holmes, O., … Grimmond, S.M. (2016). Genomic analyses identify molecular subtypes of pancreatic cancer. Nature, 531(7592), 47–52.

5. Bassett, A. R., Akhtar, A., Barlow, D. P., Bird, A. P., Brockdorff, N., Duboule, D., Ephrussi, A., Ferguson-Smith, A. C., Gingeras, T. R., Haerty, W., Higgs, D. R., Miska, E. A., & Ponting, C. P. (2014). Considerations when investigating lncRNA function in vivo. eLife, 3, e03058.

6. Brannan, C. I., Dees, E. C., Ingram, R. S., & Tilghman, S. M. (1990). The product of the H19 gene may function as an RNA. Molecular and Cellular Biology, 10(1), 28–36.

7. Brown, C. J., Ballabio, A., Rupert, J. L., Lafreniere, R. G., Grompe, M., Tonlorenzi, R., & Willard, H. F. (1991). A gene from the region of the human X inactivation centre is expressed exclusively from the inactive X chromosome. Nature, 349(6304), 38–44.

8. Cai, L., Chang, H., Fang, Y., & Li, G. (2016). A Comprehensive Characterization of the Function of LincRNAs in Transcriptional Regulation Through Long-Range Chromatin Interactions. Scientific Reports, 6, 36572.

9. Cancer Genome Atlas Research Network. (2017). Integrated Genomic Characterization of Pancreatic Ductal Adenocarcinoma. Cancer Cell, 32(2), 185–203.e13.

10. Chen, B., Dodge, M. E., Tang, W., Lu, J., Ma, Z., Fan, C.-W., Wei, S., Hao, W., Kilgore, J., Williams, N. S., Roth, M. G., Amatruda, J. F., Chen, C., & Lum, L. (2009). Small molecule-mediated disruption of Wnt-dependent signaling in tissue regeneration and cancer. Nature Chemical Biology, 5(2), 100–107.

11. Chen, J., Shishkin, A. A., Zhu, X., Kadri, S., Maza, I., Guttman, M., Hanna, J. H., Regev, A., & Garber, M. (2016). Evolutionary analysis across mammals reveals distinct classes of long non-coding RNAs. Genome Biology, 17, 19.

12. Chen, X.-F., Zhu, D.-L., Yang, M., Hu, W.-X., Duan, Y.-Y., Lu, B.-J., Rong, Y., Dong, S.-S., Hao, R.-H., Chen, J.-B., Chen, Y.-X., Yao, S., Thynn, H. N., Guo, Y., & Yang, T.-L. (2018). An Osteoporosis Risk SNP at 1p36.12 Acts as an Allele-Specific Enhancer to Modulate LINC00339 Expression via Long-Range Loop Formation. American Journal of Human Genetics, 102(5), 776–793.

13. Consortium, G., & GTEx Consortium. (2017). Genetic effects on gene expression across human tissues. Nature, 550(7675), 204–213.

14. Conway, T., Wazny, J., Bromage, A., Tymms, M., Sooraj, D., Williams, E. D., & Beresford Smith, B. (2012). Xenome--a tool for classifying reads from xenograft samples. Bioinformatics, 28(12), i172–i178.

15. Dai, L., Niu, J., & Feng, Y. (2020). Knockdown of long non coding RNA LINC00176 suppresses ovarian cancer progression by BCL3 mediated down regulation of ceruloplasmin. Journal of Cellular and Molecular Medicine, 24(1), 202–213.

16. Dixon, J. R., Selvaraj, S., Yue, F., Kim, A., Li, Y., Shen, Y., Hu, M., Liu, J. S., & Ren, B. (2012). Topological domains in mammalian genomes identified by analysis of chromatin interactions. Nature, 485(7398), 376–380.

17. Engreitz, J. M., Haines, J. E., Perez, E. M., Munson, G., Chen, J., Kane, M., McDonel, P. E., Guttman, M., & Lander, E. S. (2016). Local regulation of gene expression by lncRNA promoters, transcription and splicing. Nature, 539(7629), 452–455.

18. Esposito, R., Bosch, N., Lanzós, A., Polidori, T., Pulido-Quetglas, C., & Johnson, R. (2019). Hacking the Cancer Genome: Profiling Therapeutically Actionable Long Non-coding RNAs Using CRISPR-Cas9 Screening. Cancer Cell, 35(4), 545–557.

19. Gabay, M., Li, Y., & Felsher, D. W. (2014). MYC activation is a hallmark of cancer initiation and maintenance. Cold Spring Harbor Perspectives in Medicine, 4(6). https://doi.org/10.1101/cshperspect.a014241

20. Gao, P., Xia, J.-H., Sipeky, C., Dong, X.-M., Zhang, Q., Yang, Y., Zhang, P., Cruz, S. P., Zhang, K., Zhu, J., Lee, H.-M., Suleman, S., Giannareas, N., Liu, S., PRACTICAL Consortium, Tammela, T. L. J., Auvinen, A., Wang, X., Huang, Q., … Wei, G.-H. (2018). Biology and Clinical Implications of the 19q13 Aggressive Prostate Cancer Susceptibility Locus. Cell, 174(3), 576–589.e18.

21. Giannakis, M., Hodis, E., Jasmine Mu, X., Yamauchi, M., Rosenbluh, J., Cibulskis, K., Saksena, G., Lawrence, M. S., Qian, Z. R., Nishihara, R., Van Allen, E. M., Hahn, W. C., Gabriel, S. B., Lander, E. S., Getz, G., Ogino, S., Fuchs, C. S., & Garraway, L. A. (2014). RNF43 is frequently mutated in colorectal and endometrial cancers. Nature Genetics, 46(12), 1264– 1266.

22. Gilbert, L. A., Horlbeck, M. A., Adamson, B., Villalta, J. E., Chen, Y., Whitehead, E. H., Guimaraes, C., Panning, B., Ploegh, H. L., Bassik, M. C., Qi, L. S., Kampmann, M., & Weissman, J. S. (2014). Genome-Scale CRISPR-Mediated Control of Gene Repression and Activation. Cell, 159(3), 647–661.

23. Gilbert, L. A., Larson, M. H., Morsut, L., Liu, Z., Brar, G. A., Torres, S. E., Stern-Ginossar, N., Brandman, O., Whitehead, E. H., Doudna, J. A., Lim, W. A., Weissman, J. S., & Qi, L. S. (2013). CRISPR-mediated modular RNA-guided regulation of transcription in eukaryotes. Cell, 154(2), 442–451.

24. Gil, N., & Ulitsky, I. (2019). Regulation of gene expression by cis-acting long non-coding RNAs. Nature Reviews. Genetics. https://doi.org/10.1038/s41576-019-0184-5

25. Goldman, M., Craft, B., Hastie, M., Repečka, K., McDade, F., Kamath, A., Banerjee, A., Luo, Y., Rogers, D., Brooks, A. N., Zhu, J., & Haussler, D. (2019). The UCSC Xena platform for public and private cancer genomics data visualization and interpretation. bioRxiv, 326470. doi: https://doi.org/10.1101/326470

26. Goudarzi, M., Berg, K., Pieper, L. M., & Schier, A. F. (2019). Individual long non-coding RNAs have no overt functions in zebrafish embryogenesis, viability and fertility. eLife, 8. https://doi.org/10.7554/eLife.40815

27. Guttman, M., Donaghey, J., Carey, B. W., Garber, M., Grenier, J. K., Munson, G., Young, G., Lucas, A. B., Ach, R., Bruhn, L., Yang, X., Amit, I., Meissner, A., Regev, A., Rinn, J. L., Root, D. E., & Lander, E. S. (2011). lincRNAs act in the circuitry controlling pluripotency and differentiation. Nature, 477(7364), 295–300.

28. Han, X., Luo, S., Peng, G., Lu, J. Y., Cui, G., Liu, L., Yan, P., Yin, Y., Liu, W., Wang, R., Zhang, J., Ai, S., Chang, Z., Na, J., He, A., Jing, N., & Shen, X. (2018). Mouse knockout models reveal largely dispensable but context-dependent functions of lncRNAs during development. Journal of Molecular Cell Biology, 10(2), 175–178.

29. Hensman, J., Lawrence, N. D., & Rattray, M. (2013). Hierarchical Bayesian modelling of gene expression time series across irregularly sampled replicates and clusters. BMC Bioinformatics, 14, 252.

30. Hindorff, L. A., Sethupathy, P., Junkins, H. A., Ramos, E. M., Mehta, J. P., Collins, F. S., & Manolio, T. A. (2009). Potential etiologic and functional implications of genome-wide association loci for human diseases and traits. Proceedings of the National Academy of Sciences of the United States of America, 106(23), 9362–9367.

31. Holtzhausen, A., Zhao, F., Evans, K. S., Tsutsui, M., Orabona, C., Tyler, D. S., & Hanks, B. A. (2015). Melanoma-Derived Wnt5a Promotes Local Dendritic-Cell Expression of IDO and Immunotolerance: Opportunities for Pharmacologic Enhancement of Immunotherapy. Cancer Immunology Research, 3(9), 1082–1095.

32. Hon, C.-C., Ramilowski, J. A., Harshbarger, J., Bertin, N., Rackham, O. J. L., Gough, J., Denisenko, E., Schmeier, S., Poulsen, T. M., Severin, J., Lizio, M., Kawaji, H., Kasukawa, T., Itoh, M., Burroughs, A. M., Noma, S., Djebali, S., Alam, T., Medvedeva, Y. A., … Forrest, A. R. R. (2017). An atlas of human long non-coding RNAs with accurate 5’ ends. Nature, 543(7644), 199–204.

33. Horlbeck, M. A., Gilbert, L. A., Villalta, J. E., Adamson, B., Pak, R. A., Chen, Y., Fields, A. P., Park, C. Y., Corn, J. E., Kampmann, M., & Weissman, J. S. (2016). Compact and highly active next-generation libraries for CRISPR-mediated gene repression and activation. eLife, 5. https://doi.org/10.7554/eLife.19760

34. Huarte, M. (2015). The emerging role of lncRNAs in cancer. Nature Medicine, 21(11), 1253– 1261.

35. Huarte, M., Guttman, M., Feldser, D., Garber, M., Koziol, M. J., Kenzelmann-Broz, D., Khalil, A. M., Zuk, O., Amit, I., Rabani, M., Attardi, L. D., Regev, A., Lander, E. S., Jacks, T., & Rinn, J. L. (2010). A Large Intergenic Noncoding RNA Induced by p53 Mediates Global Gene Repression in the p53 Response. Cell, 142(3), 409–419.

36. Idris, M., Harmston, N., Petretto, E., Madan, B., & Virshup, D. M. (2019). Broad regulation of gene isoform expression by Wnt signaling in cancer. RNA, 25(12), 1696–1713.

37. Iyer, M. K., Niknafs, Y. S., Malik, R., Singhal, U., Sahu, A., Hosono, Y., Barrette, T. R., Prensner, J. R., Evans, J. R., Zhao, S., Poliakov, A., Cao, X., Dhanasekaran, S. M., Wu, Y.-M., Robinson, D. R., Beer, D. G., Feng, F. Y., Iyer, H. K., & Chinnaiyan, A. M. (2015). The landscape of long noncoding RNAs in the human transcriptome. Nature Genetics, 47(3), 199–208.

38. Jiang, X., Hao, H.-X., Growney, J. D., Woolfenden, S., Bottiglio, C., Ng, N., Lu, B., Hsieh, M. H., Bagdasarian, L., Meyer, R., Smith, T. R., Avello, M., Charlat, O., Xie, Y., Porter, J. A., Pan, S., Liu, J., McLaughlin, M. E., & Cong, F. (2013). Inactivating mutations of RNF43 confer Wnt dependency in pancreatic ductal adenocarcinoma. Proceedings of the National Academy of Sciences of the United States of America, 110(31), 12649–12654.

39. Jiang, Y., Jiang, Y.-Y., Xie, J.-J., Mayakonda, A., Hazawa, M., Chen, L., Xiao, J.-F., Li, C.-Q., Huang, M.-L., Ding, L.-W., Sun, Q.-Y., Xu, L., Kanojia, D., Jeitany, M., Deng, J.-W., Liao, L.-D., Soukiasian, H. J., Berman, B. P., Hao, J.-J., … Koeffler, H. P. (2018). Co-activation of super-enhancer-driven CCAT1 by TP63 and SOX2 promotes squamous cancer progression. Nature Communications, 9(1), 3619.

40. Joung, J., Engreitz, J. M., Konermann, S., Abudayyeh, O. O., Verdine, V. K., Aguet, F., Gootenberg, J. S., Sanjana, N. E., Wright, J. B., Fulco, C. P., Tseng, Y.-Y., Yoon, C. H., Boehm, J. S., Lander, E. S., & Zhang, F. (2017). Genome-scale activation screen identifies a lncRNA locus regulating a gene neighbourhood. Nature, 548(7667), 343–346.

41. Kawasaki, Y., Komiya, M., Matsumura, K., Negishi, L., Suda, S., Okuno, M., Yokota, N., Osada, T., Nagashima, T., Hiyoshi, M., Okada-Hatakeyama, M., Kitayama, J., Shirahige, K., & Akiyama, T. (2016). MYU, a Target lncRNA for Wnt/c-Myc Signaling, Mediates Induction of CDK6 to Promote Cell Cycle Progression. Cell Reports, 16(10), 2554–2564.

42. Killion, J. J., Radinsky, R., & Fidler, I. J. (1998). Orthotopic models are necessary to predict therapy of transplantable tumors in mice. Cancer Metastasis Reviews, 17(3), 279–284.

43. Kim, D., Pertea, G., Trapnell, C., Pimentel, H., Kelley, R., & Salzberg, S. L. (2013). TopHat2: accurate alignment of transcriptomes in the presence of insertions, deletions and gene fusions. Genome Biology, 14(4), R36.

44. Kohtz, J. D. (2014). Long non-coding RNAs learn the importance of being in vivo. Frontiers in Genetics, 5, 45.

45. Kong, L., Zhang, Y., Ye, Z.-Q., Liu, X.-Q., Zhao, S.-Q., Wei, L., & Gao, G. (2007). CPC: assess the protein-coding potential of transcripts using sequence features and support vector machine. Nucleic Acids Research, 35(Web Server issue), W345–W349.

46. Koo, B.-K., Spit, M., Jordens, I., Low, T. Y., Stange, D. E., van de Wetering, M., van Es, J. H., Mohammed, S., Heck, A. J. R., Maurice, M. M., & Clevers, H. (2012). Tumour suppressor RNF43 is a stem-cell E3 ligase that induces endocytosis of Wnt receptors. Nature, 488(7413), 665–669.

47. Kopp, F., & Mendell, J. T. (2018). Functional Classification and Experimental Dissection of Long Noncoding RNAs. Cell, 172(3), 393–407.

48. Krishnan, V. (2006). Regulation of bone mass by Wnt signaling. Journal of Clinical Investigation, 116(5), 1202–1209.

49. Letai, A. (2017). Functional precision cancer medicine—moving beyond pure genomics. Nature Medicine, 23(9), 1028–1035.

50. Li, Q., Seo, J.-H., Stranger, B., McKenna, A., Pe’er, I., Laframboise, T., Brown, M., Tyekucheva, S., & Freedman, M. L. (2013). Integrative eQTL-based analyses reveal the biology of breast cancer risk loci. Cell, 152(3), 633–641.

51. Liu, S. J., Horlbeck, M. A., Cho, S. W., Birk, H. S., Malatesta, M., He, D., Attenello, F. J., Villalta, J. E., Cho, M. Y., Chen, Y., Mandegar, M. A., Olvera, M. P., Gilbert, L. A., Conklin, B. R., Chang, H. Y., Weissman, J. S., & Lim, D. A. (2017). CRISPRi-based genome-scale identification of functional long noncoding RNA loci in human cells. Science, 355(6320). https://doi.org/10.1126/science.aah7111

52. Liu, Y., Cao, Z., Wang, Y., Guo, Y., Xu, P., Yuan, P., Liu, Z., He, Y., & Wei, W. (2018). Genome-wide screening for functional long noncoding RNAs in human cells by Cas9 targeting of splice sites. Nature Biotechnology. https://doi.org/10.1038/nbt.4283

53. Li, W., Xu, H., Xiao, T., Cong, L., Love, M. I., Zhang, F., Irizarry, R. A., Liu, J. S., Brown, M., & Liu, X. S. (2014). MAGeCK enables robust identification of essential genes from genome-scale CRISPR/Cas9 knockout screens. Genome Biology, 15(12), 554.

54. Love, M. I., Huber, W., & Anders, S. (2014). Moderated estimation of fold change and dispersion for RNA-seq data with DESeq2. Genome Biology, 15(12), 550.

55. Luo, S., Lu, J. Y., Liu, L., Yin, Y., Chen, C., Han, X., Wu, B., Xu, R., Liu, W., Yan, P., Shao, W., Lu, Z., Li, H., Na, J., Tang, F., Wang, J., Zhang, Y. E., & Shen, X. (2016). Divergent lncRNAs Regulate Gene Expression and Lineage Differentiation in Pluripotent Cells. Cell Stem Cell, 18(5), 637–652.

56. Madan, B., Harmston, N., Nallan, G., Montoya, A., Faull, P., Petretto, E., & Virshup, D. M. (2018). Temporal dynamics of Wnt-dependent transcriptome reveal an oncogenic Wnt/MYC/ribosome axis. The Journal of Clinical Investigation, 128(12), 5620–5633.

57. Madan, B., Ke, Z., Harmston, N., Ho, S. Y., Frois, A. O., Alam, J., Jeyaraj, D. A., Pendharkar, V., Ghosh, K., Virshup, I. H., Manoharan, V., Ong, E. H. Q., Sangthongpitag, K., Hill, J., Petretto, E., Keller, T. H., Lee, M. A., Matter, A., & Virshup, D. M. (2016). Wnt addiction of genetically defined cancers reversed by PORCN inhibition. Oncogene, 35(17), 2197–2207.

58. Madan, B., & Virshup, D. M. (2015). Targeting Wnts at the Source--New Mechanisms, New Biomarkers, New Drugs. Molecular Cancer Therapeutics, 14(5), 1087–1094.

59. Miller, T. E., Liau, B. B., Wallace, L. C., Morton, A. R., Xie, Q., Dixit, D., Factor, D. C., Kim, L. J. Y., Morrow, J. J., Wu, Q., Mack, S. C., Hubert, C. G., Gillespie, S. M., Flavahan, W. A., Hoffmann, T., Thummalapalli, R., Hemann, M. T., Paddison, P. J., Horbinski, C. M., … Rich, J. N. (2017). Transcription elongation factors represent in vivo cancer dependencies in glioblastoma. Nature, 547(7663), 355–359.

60. Muir, A., & Vander Heiden, M. G. (2018). The nutrient environment affects therapy. Science, 360(6392), 962–963.

61. Ng, M., Tan, D. S. P., Subbiah, V., Weekes, C. D., Teneggi, V., Diermayr, V., Ethirajulu, K., Yeo, P., Chen, D., Blanchard, S., Nellore, R., Gan, B. H., Yasin, M., Lee, L. H., Lee, M. A., Hill, J., Madan, B., Virshup, D., & Matter, A. (2017). First-in-human phase 1 study of ETC-159 an oral PORCN inhbitor in patients with advanced solid tumours. Journal of Clinical Oncology, 35(15_suppl), 2584–2584.

62. Nusse, R., & Clevers, H. (2017). Wnt/β-Catenin Signaling, Disease, and Emerging Therapeutic Modalities. Cell, 169(6), 985–999.

63. Polakis, P. (2012). Wnt Signaling in Cancer. Cold Spring Harbor Perspectives in Biology, 4(5), a008052–a008052.

64. Possik, P. A., Müller, J., Gerlach, C., Kenski, J. C. N., Huang, X., Shahrabi, A., Krijgsman, O., Song, J.-Y., Smit, M. A., Gerritsen, B., Lieftink, C., Kemper, K., Michaut, M., Beijersbergen, R. L., Wessels, L., Schumacher, T. N., & Peeper, D. S. (2014). Parallel in vivo and in vitro melanoma RNAi dropout screens reveal synthetic lethality between hypoxia and DNA damage response inhibition. Cell Reports, 9(4), 1375–1386.

65. Powell, J. E., Fung, J. N., Shakhbazov, K., Sapkota, Y., Cloonan, N., Hemani, G., Hillman, K. M., Kaufmann, S., Luong, H. T., Bowdler, L., Painter, J. N., Holdsworth-Carson, S. J., Visscher, P. M., Dinger, M. E., Healey, M., Nyholt, D. R., French, J. D., Edwards, S. L., Rogers, P. A. W., & Montgomery, G. W. (2016). Endometriosis risk alleles at 1p36.12 act through inverse regulation of CDC42 and LINC00339. Human Molecular Genetics, 25(22), 5046–5058.

66. Reimand, J., Arak, T., Adler, P., Kolberg, L., Reisberg, S., Peterson, H., & Vilo, J. (2016). g:Profiler—a web server for functional interpretation of gene lists (2016 update). Nucleic Acids Research, 44(W1), W83–W89.

67. Rotival, M., & Petretto, E. (2014). Leveraging gene co-expression networks to pinpoint the regulation of complex traits and disease, with a focus on cardiovascular traits. Briefings in Functional Genomics, 13(1), 66–78.

68. Ruan, X., Li, P., Chen, Y., Shi, Y., Pirooznia, M., Seifuddin, F., Suemizu, H., Ohnishi, Y., Yoneda, N., Nishiwaki, M., Shepherdson, J., Suresh, A., Singh, K., Ma, Y., Jiang, C.-F., & Cao, H. (2020). In vivo functional analysis of non-conserved human lncRNAs associated with cardiometabolic traits. Nature Communications, 11(1), 45.

69. Schmitt, A. M., & Chang, H. Y. (2016). Long Noncoding RNAs in Cancer Pathways. Cancer Cell, 29(4), 452–463.

70. Seshagiri, S., Stawiski, E. W., Durinck, S., Modrusan, Z., Storm, E. E., Conboy, C. B., Chaudhuri, S., Guan, Y., Janakiraman, V., Jaiswal, B. S., Guillory, J., Ha, C., Dijkgraaf, G. J. P., Stinson, J., Gnad, F., Huntley, M. A., Degenhardt, J. D., Haverty, P. M., Bourgon, R., … de Sauvage, F. J. (2012). Recurrent R-spondin fusions in colon cancer. Nature, 488(7413), 660–664.

71. Sharma, S. V., Haber, D. A., & Settleman, J. (2010). Cell line-based platforms to evaluate the therapeutic efficacy of candidate anticancer agents. Nature Reviews. Cancer, 10(4), 241– 253.

72. Smith, I., Greenside, P. G., Natoli, T., Lahr, D. L., Wadden, D., Tirosh, I., Narayan, R., Root, D. E., Golub, T. R., Subramanian, A., & Doench, J. G. (2017). Evaluation of RNAi and CRISPR technologies by large-scale gene expression profiling in the Connectivity Map. PLoS Biology, 15(11), e2003213.

73. Spranger, S., Bao, R., & Gajewski, T. F. (2015). Melanoma-intrinsic β-catenin signalling prevents anti-tumour immunity. Nature, 523(7559), 231–235.

74. Stojic, L., Lun, A. T. L., Mangei, J., Mascalchi, P., Quarantotti, V., Barr, A. R., Bakal, C., Marioni, J. C., Gergely, F., & Odom, D. T. (2018). Specificity of RNAi, LNA and CRISPRi as loss-of-function methods in transcriptional analysis. Nucleic Acids Research, 46(12), 5950–5966.

75. Tan, J. Y., Smith, A. A. T., Ferreira da Silva, M., Matthey-Doret, C., Rueedi, R., Sönmez, R., Ding, D., Kutalik, Z., Bergmann, S., & Marques, A.C. (2017). cis-Acting Complex-Trait-Associated lincRNA Expression Correlates with Modulation of Chromosomal Architecture. Cell Reports, 18(9), 2280–2288.

76. Tran, D. D. H., Kessler, C., Niehus, S. E., Mahnkopf, M., Koch, A., & Tamura, T. (2018). Myc target gene, long intergenic noncoding RNA, Linc00176 in hepatocellular carcinoma regulates cell cycle and cell survival by titrating tumor suppressor microRNAs. Oncogene, 37(1), 75–85.

77. Trapnell, C., Williams, B. A., Pertea, G., Mortazavi, A., Kwan, G., van Baren, M. J., Salzberg, S. L., Wold, B. J., & Pachter, L. (2010). Transcript assembly and quantification by RNA-Seq reveals unannotated transcripts and isoform switching during cell differentiation. Nature Biotechnology, 28(5), 511–515.

78. Trimarchi, T., Bilal, E., Ntziachristos, P., Fabbri, G., Dalla-Favera, R., Tsirigos, A., & Aifantis, I. (2014). Genome-wide mapping and characterization of Notch-regulated long noncoding RNAs in acute leukemia. Cell, 158(3), 593–606.

79. Waddell, N., Pajic, M., Patch, A.-M., Chang, D. K., Kassahn, K. S., Bailey, P., Johns, A. L., Miller, D., Nones, K., Quek, K., Quinn, M. C. J., Robertson, A. J., Fadlullah, M. Z. H., Bruxner, T. J. C., Christ, A. N., Harliwong, I., Idrisoglu, S., Manning, S., Nourse, C., … Grimmond, S. M. (2015). Whole genomes redefine the mutational landscape of pancreatic cancer. Nature, 518(7540), 495–501.

80. Wang, L., Park, H. J., Dasari, S., Wang, S., Kocher, J.-P., & Li, W. (2013). CPAT: Coding-Potential Assessment Tool using an alignment-free logistic regression model. Nucleic Acids Research, 41(6), e74.

81. Wang, Y., Hanifi-Moghaddam, P., Hanekamp, E. E., Kloosterboer, H. J., Franken, P., Veldscholte, J., van Doorn, H. C., Ewing, P. C., Kim, J. J., Grootegoed, J. A., Burger, C. W., Fodde, R., & Blok, L. J. (2009). Progesterone inhibition of Wnt/beta-catenin signaling in normal endometrium and endometrial cancer. Clinical Cancer Research, 15(18), 5784– 5793.

82. Wang, Y., Song, F., Zhang, B., Zhang, L., Xu, J., Kuang, D., Li, D., Choudhary, M. N. K., Li, Y., Hu, M., Hardison, R., Wang, T., & Yue, F. (2018). The 3D Genome Browser: a web-based browser for visualizing 3D genome organization and long-range chromatin interactions. Genome Biology, 19(1), 151.

83. Whiteside, T. L. (2008). The tumor microenvironment and its role in promoting tumor growth. Oncogene, 27(45), 5904–5912.

84. Willert, K., Brown, J. D., Danenberg, E., Duncan, A. W., Weissman, I. L., Reya, T., Yates, J. R., 3rd, & Nusse, R. (2003). Wnt proteins are lipid-modified and can act as stem cell growth factors. Nature, 423(6938), 448–452.

85. Xiang, J.-F., Yin, Q.-F., Chen, T., Zhang, Y., Zhang, X.-O., Wu, Z., Zhang, S., Wang, H.-B., Ge, J., Lu, X., Yang, L., & Chen, L.-L. (2014). Human colorectal cancer-specific CCAT1-L lncRNA regulates long-range chromatin interactions at the MYC locus. Cell Research, 24(5), 513–531.

86. Yau, E. H., Kummetha, I. R., Lichinchi, G., Tang, R., Zhang, Y., & Rana, T. M. (2017). Genome-Wide CRISPR Screen for Essential Cell Growth Mediators in Mutant KRAS Colorectal Cancers. Cancer Research, 77(22), 6330–6339.

87. Yuan, J.-H., Yang, F., Wang, F., Ma, J.-Z., Guo, Y.-J., Tao, Q.-F., Liu, F., Pan, W., Wang, T.-T., Zhou, C.-C., Wang, S.-B., Wang, Y.-Z., Yang, Y., Yang, N., Zhou, W.-P., Yang, G.-S., & Sun, S.-H. (2014). A long noncoding RNA activated by TGF-β promotes the invasion-metastasis cascade in hepatocellular carcinoma. Cancer Cell, 25(5), 666–681.

88. Zhang, E., Han, L., Yin, D., He, X., Hong, L., Si, X., Qiu, M., Xu, T., De, W., Xu, L., Shu, Y., & Chen, J. (2017). H3K27 acetylation activated-long non-coding RNA CCAT1 affects cell proliferation and migration by regulating SPRY4 and HOXB13 expression in esophageal squamous cell carcinoma. Nucleic Acids Research, 45(6), 3086–3101.

89. Zhang, J., Sui, S., Wu, H., Zhang, J., Zhang, X., Xu, S., & Pang, D. (2019). The transcriptional landscape of lncRNAs reveals the oncogenic function of LINC00511 in ER-negative breast cancer. Cell Death & Disease, 10(8), 599.

90. Zhan, T., Rindtorff, N., & Boutros, M. (2017). Wnt signaling in cancer. Oncogene, 36(11), 1461– 1473.

91. Zhong, Z., Sepramaniam, S., Chew, X. H., Wood, K., Lee, M. A., Madan, B., & Virshup, D. M. (2019). PORCN inhibition synergizes with PI3K/mTOR inhibition in Wnt-addicted cancers. Oncogene, 38(40), 6662–6677.

92. Zhong, Z., & Virshup, D. M. (2019). Wnt signaling and drug resistance in cancer. Molecular Pharmacology. https://doi.org/10.1124/mol.119.117978

93. Zhu, S., Li, W., Liu, J., Chen, C.-H., Liao, Q., Xu, P., Xu, H., Xiao, T., Cao, Z., Peng, J., Yuan, P., Brown, M., Liu, X. S., & Wei, W. (2016). Genome-scale deletion screening of human long non-coding RNAs using a paired-guide RNA CRISPR–Cas9 library. Nature Biotechnology, 34(12), 1279–1286.

