## Supplementary material for "Wnt-regulated lncRNA discovery enhanced by *in vivo* identification and CRISPRi functional validation": Figure S1

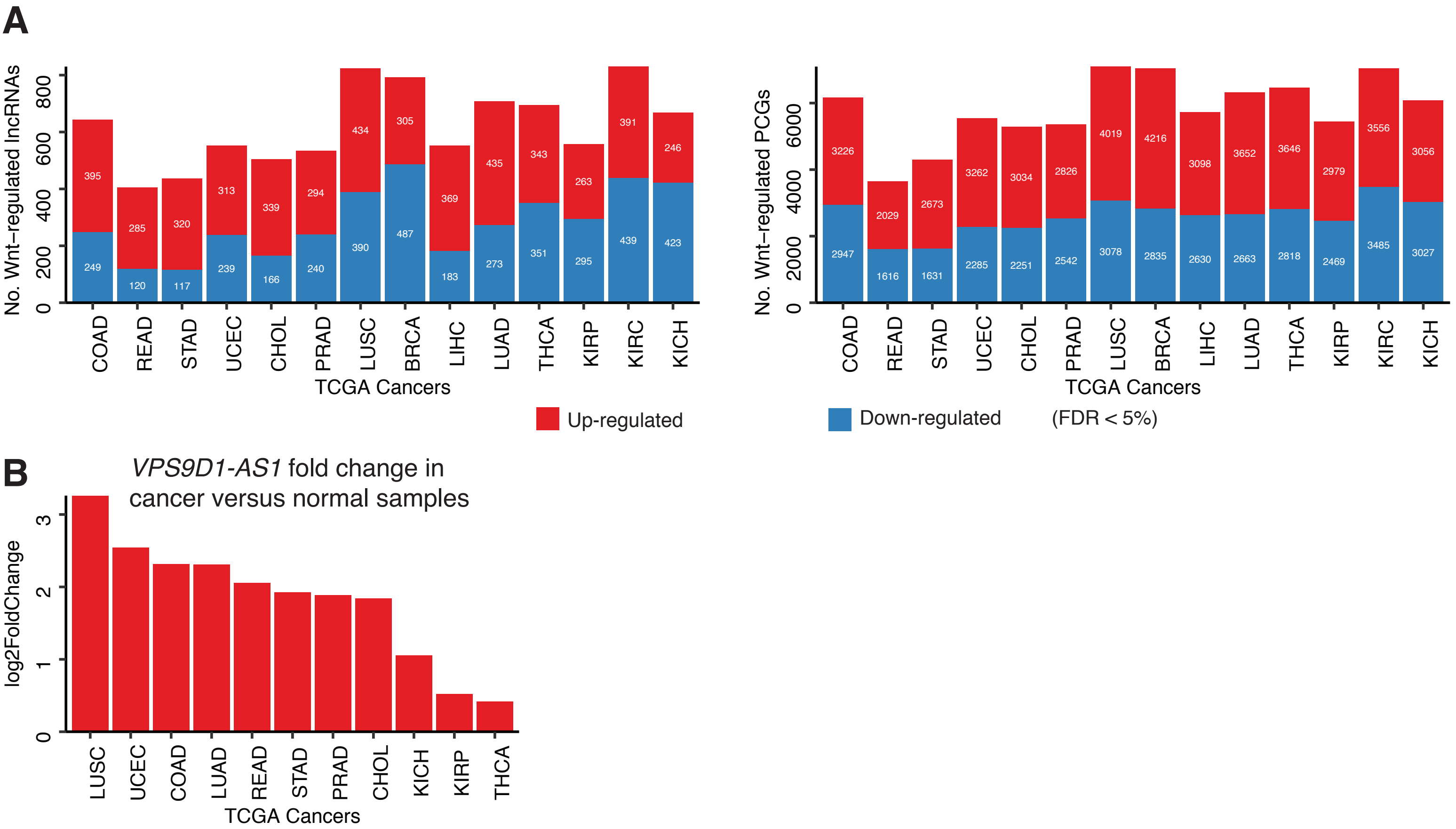


**Figure S1. Wnt-regulated lncRNAs and PCGs are dysregulated in TCGA cancers (A) Wnt-regulated lncRNAs and PCGs, defined as genes changed over time upon Wnt inhibition (FDR < 5%) in the orthotopic *RNF43*-mutant pancreatic cancer model (Figure 1A). Wnt-regulated lncRNAs and PCGs are dysregulated in different types of cancers as determined by differential expression between tumors and their paired normal samples using the TCGA dataset. (B) *VPS9D-AS1* is upregulated in 11 different types of cancers.**
