## Supplementary material for "Wnt-regulated lncRNA discovery enhanced by *in vivo* identification and CRISPRi functional validation": Figure S2

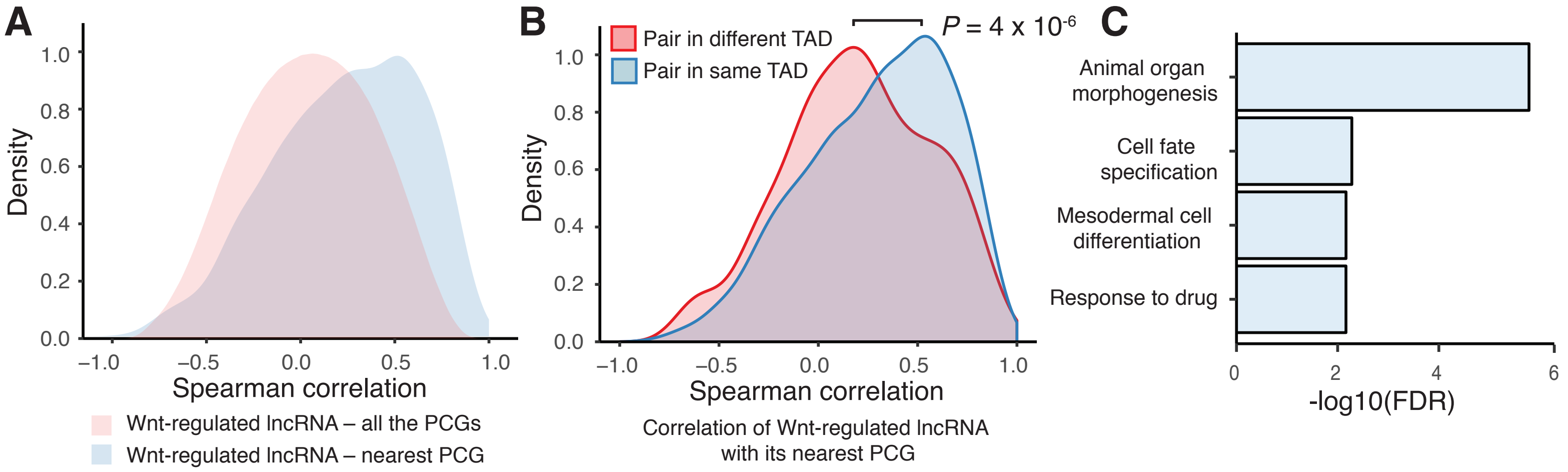


**Figure S2. Subset of Wnt-dependent lncRNAs co-express with its nearest PCG in the same TAD. (A) Wnt-regulated lncRNAs exhibit stronger co-expression with their nearest PCG after Wnt inhibition compared to their co-expression with all PCGs. (B) Wnt-regulated lncRNA–nearest PCG pairs within the same TAD exhibit stronger co-expression than the pairs in different TADs. *P* for significance was calculated by Mann–Whitney U test. (C) For the Wnt-regulated lncRNA–nearest PCG pairs encoded within the same TAD, the PCGs are significantly (FDR < 5%) enriched for GO biological processes.**
