## Supplementary material for "Wnt-regulated lncRNA discovery enhanced by *in vivo* identification and CRISPRi functional validation": Figure S3

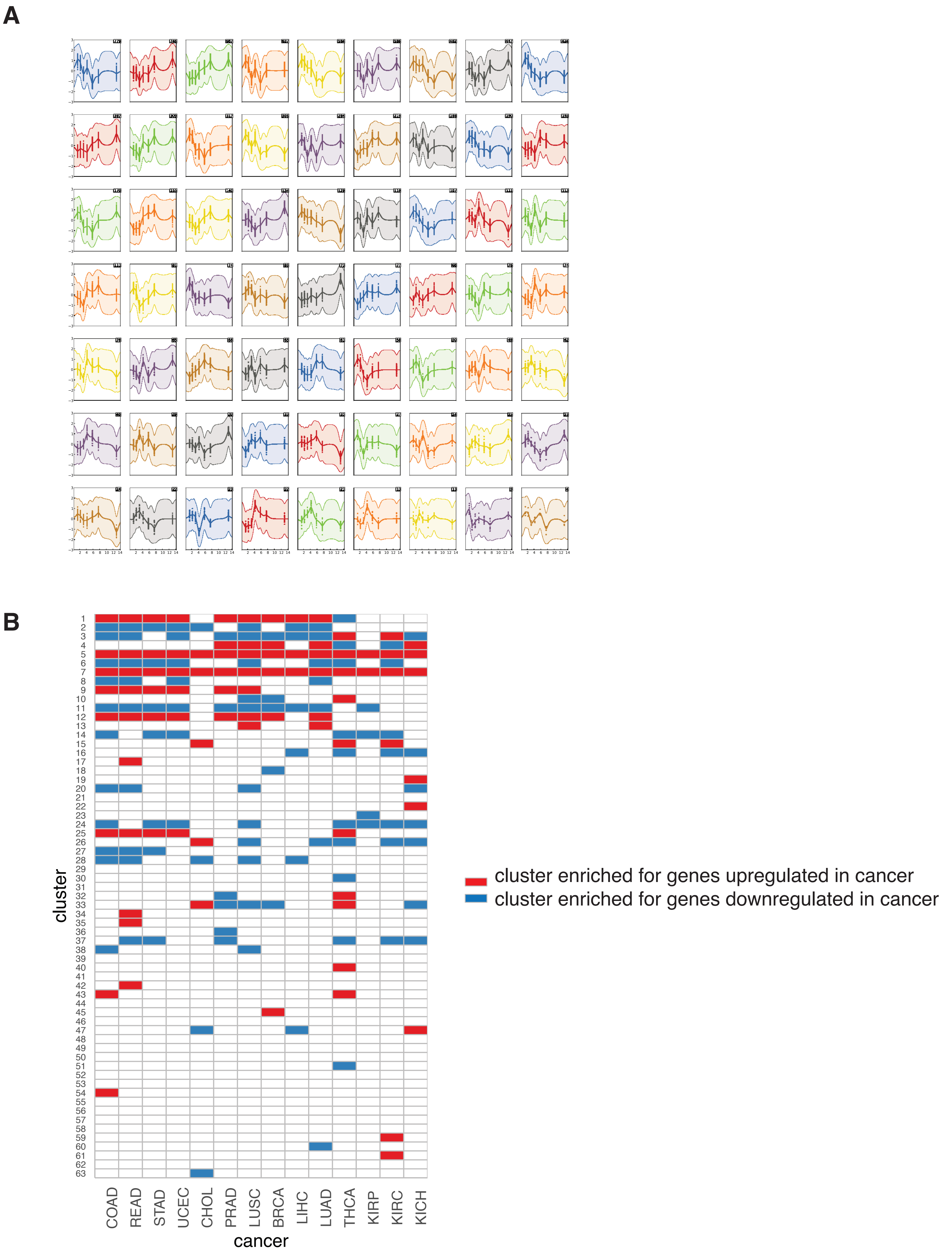


**Figure S3. Clusters are enriched for genes dysregulated in different cancers. (A) The Wnt-regulated lncRNAs and PCGs fall into 63 distinct clusters based on their pattern of expression change following Wnt inhibition. (B) 46 out of the 63 clusters are enriched (FDR < 5%) for genes dysregulated in at least one type of cancer.**
