## Supplementary material for "Wnt-regulated lncRNA discovery enhanced by *in vivo* identification and CRISPRi functional validation": Figure S4

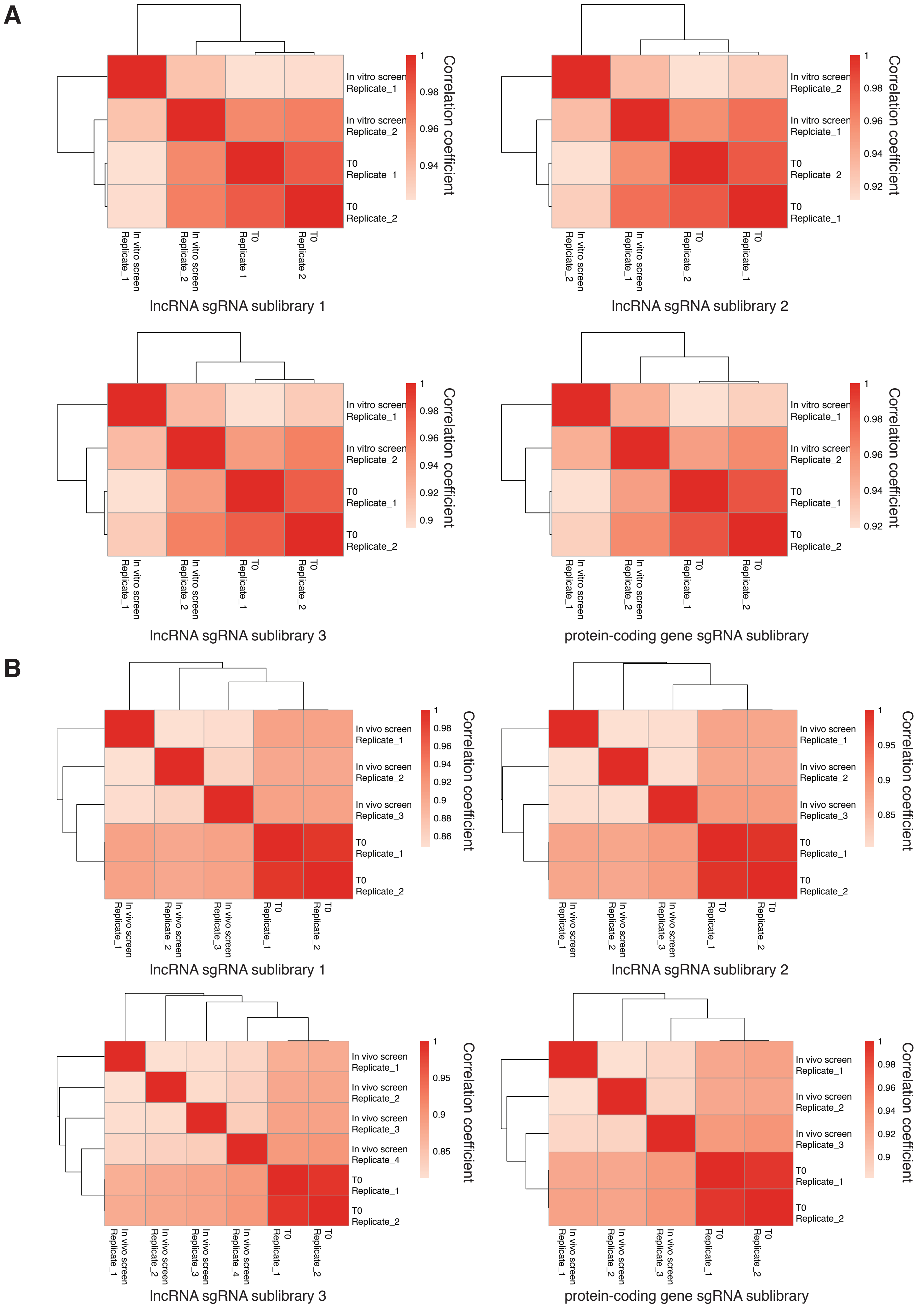


**Figure S4. A high correlation of sgRNA counts between independent experimental replicates in CRISPRi screens. (A) Correlation of sgRNA counts between experimental replicates in the *in vitro* screens. (B) Correlation of sgRNA counts between experimental replicates in the *in vivo* screens.**
