## Supplementary figures and images for "Wnt-regulated lncRNA discovery enhanced by *in vivo* identification and CRISPRi functional validation"

### Figure S5

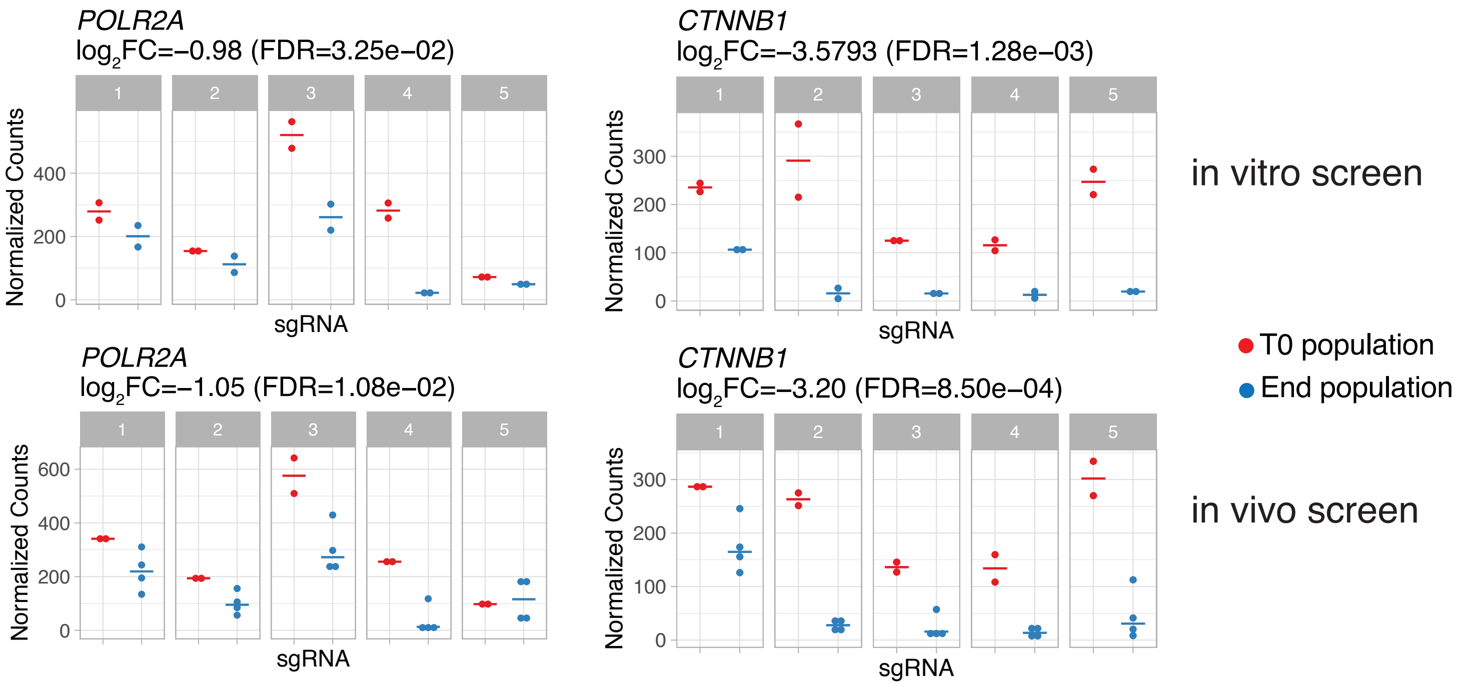


**Figure S5. CRISPRi screens are able to identify important positive controls as gene hits.**

### Figure S6

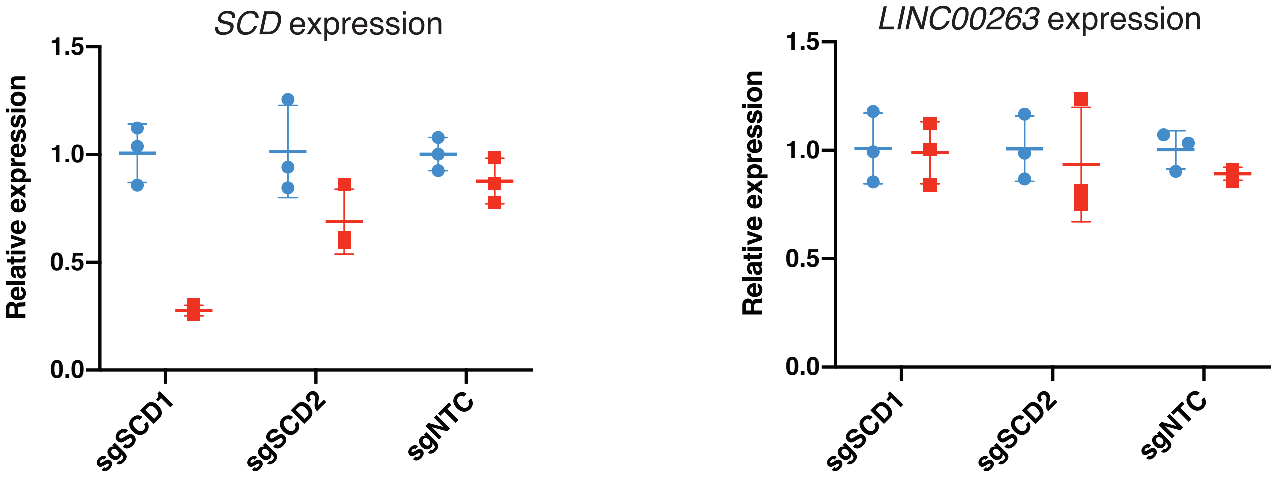


**Figure S6. Knockdown of *SCD* with CRISPRi reduce *SCD* mRNA abundance, but not the expression of *LINC00263*.**
